# Dynamic interplay between target search and recognition for the Cascade surveillance complex of type I-E CRISPR-Cas systems

**DOI:** 10.1101/2022.12.18.520913

**Authors:** Pierre Aldag, Marius Rutkauskas, Julene Madariaga-Marcos, Inga Songailiene, Tomas Sinkunas, Felix E Kemmerich, Dominik J Kauert, Virginijus Siksnys, Ralf Seidel

## Abstract

CRISPR-Cas effector complexes enable the defense against foreign nucleic acids and have recently been exploited as molecular tools for precise genome editing at a target locus. To bind and cleave their target, the CRISPR-Cas effectors first have to interrogate the entire genome for the presence of a matching sequence. Matching is achieved by base-pairing between the crRNA of the complexes and the DNA target strand such that an R-loop is formed. R-loop formation starts at a specific PAM motif and progresses reversibly in single base-pair steps until mismatches stop further progression or until the full target is recognized and destroyed. The reversible nature of this process entails that even a fully matching target should only become recognized with a low probability per target encounter. The details of this process, which directly affect the effectiveness of the target search, remain unresolved.

Here we dissect the target search process of the Type I CRISPR-Cas complex Cascade by simultaneously monitoring DNA binding and R-loop formation by the complex. We directly quantify the low target recognition probabilities and show that they increase with increasing negative supercoiling. Furthermore, we demonstrate that Cascade uses a combination of three-dimensional and limited one-dimensional diffusion along the DNA contour for its target search. The latter allows for rapidly scanning the PAM sequences in a given region and, importantly, significantly increasing the overall efficiency of the target search by repeatedly revisiting the sites. Overall we show that target search and target recognition are tightly linked and that DNA supercoiling and limited 1D diffusion need to be considered when understanding target recognition and target search by CRISPR-Cas enzymes and engineering more efficient and precise variants.

## Introduction

CRISPR (Clustered Regularly Interspaced Palindromic Repeats)-Cas (CRISPR associated) systems constitute adaptive immune systems in prokaryotes^1–3^. The Cascade surveillance complex of the type I-E CRISPR-Cas system of *Streptococcus thermophilus* (St-Cascade thereafter) is a large multi-subunit RNA-guided ribonucleoprotein complex of ∼400 kDa molecular weight, which site-specifically targets double-stranded DNA^4–7^. Type I CRISPR-Cas surveillance complexes have shown promising potential for applications in genome editing^8–11^. A prerequisite for target recognition is the binding of the surveillance complexes to a short sequence called protospacer adjacent motif (PAM)^12,13^. PAM recognition triggers an initial melting of the adjacent DNA duplex and primes base pairing between the guide RNA and the complementary target strand while displacing the non-target strand (Fig. 1a). The resulting structure is called an R-loop^14,15^. R-loop formation is achieved by unwinding the DNA from the PAM towards the PAM-distal end of the target in a reversible process driven by thermal fluctuations^16–18^. For St-Cascade, the R-loop is 32 base pairs (bp) long and the PAM corresponds to an AAN motif. However, St-Cascade also tolerates other dinucleotide PAM combinations than AA, albeit with lower binding affinities^4^. Once a complete R-loop has been formed, Cascade undergoes a conformational change that stably ‘locks’ the protein complex on the DNA^6,15,19^. This allows recruitment of the ATP-driven nuclease-helicase Cas3, which then degrades the non-target strand in a 3’-to-5’ direction^20–24^. Mismatches between the guide RNA and the target strand hinder R-loop expansion beyond the mismatch position, hence promoting R-loop collapse^16^. PAM-proximal mismatches typically impede the R-loop progression more strongly than PAM-distal mismatches^25,26^. Once mismatches are overcome, R-loops become locked and cleaved similarly to the wild-type target^16^. However, more than five continuous PAM-distal mismatches abolish ‘locking’ and, thus, cleavage^15,16^.

**Figure 1.**
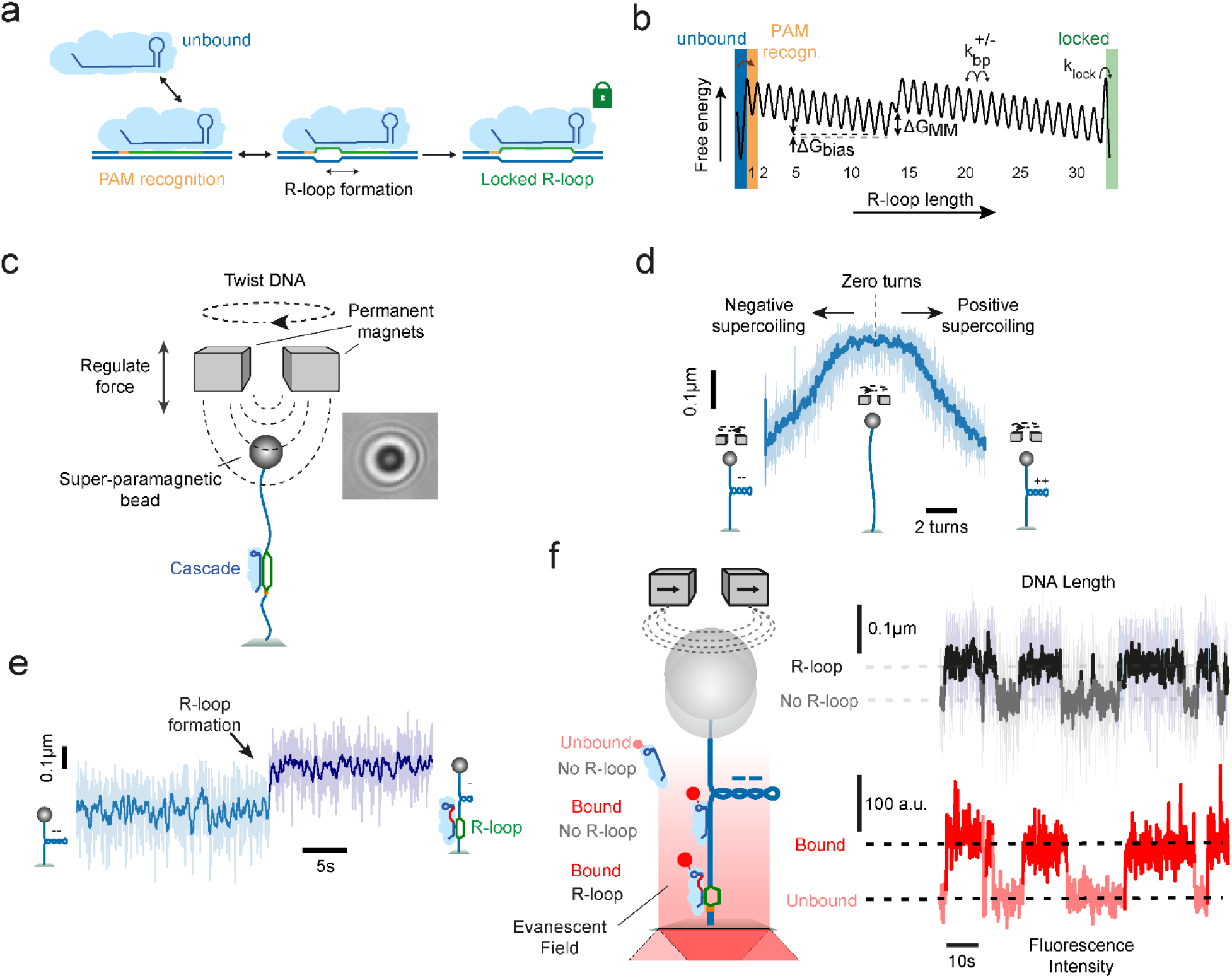
Experimental setup to detect DNA binding and R-loop formation by St-Cascade. **a** Scheme of target recognition by St-Cascade including PAM binding, R-loop formation and stable locking once the R-loop becomes fully extended. **b** Simplified 1D energy landscape for R-loop formation by St-Cascade. For states 1 to 32, the index corresponds to the R-loop length. Unbound, PAM-bound and locked state are indicated by colored bars. Negative supercoiling introduces a constant negative bias of Δ*G*_*bias*_ per bp. A mismatch between the target strand and the guide RNA introduces a local free energy penalty Δ*G*_*MM*_. **c** (left) Scheme of the magnetic tweezers setup used to stretch and twist single double-stranded DNA molecules. (right) Observed diffraction pattern of a ∼1 µm magnetic bead, used to track the length of the DNA molecule in real time. **d** DNA supercoiling curve taken at 0.3 pN, showing the characteristic DNA length reduction due to writhe formation upon negative and positive supercoiling. **e** Time trajectory of the DNA length including an R-loop formation event by St-Cascade seen as a sudden length increase, shown here for an R-loop length of 12bp. Data was taken at -6 turns and 0.3 pN. **f** (left) Scheme of the combined magnetic tweezers and TIRF microscopy measurements. Illuminating the DNA near its surface attachment with an evanescent field additionally allows to detect binding and dissociation of St-Cascade. (right) Correlated time trajectories of the DNA length monitoring R-loop formation and of the fluorescent intensity monitoring Cascade binding. In all subpanels, the shown DNA length data was taken at 120 Hz (light colors) and smoothed to 3 Hz (dark colors), while the shown fluorescence data was taken at 10 Hz.

The reversible R-loop formation process is thought to be the actual target discrimination mechanism^16,27^. Any potential target with a permissive PAM in a genome gets explored until mismatches hinder R-loop expansion and promote collapse. A rapid reversibility of R-loop formation, thus, circumvents stalling of the protein complex at partially matching sites to allow rapid scanning of all potential sites during target search. Reversible R-loop formation, and similarly other strand exchange reactions, have been previously modelled as a random walk of 1 bp steps within a simplified one-dimensional energy landscape, starting at the PAM and ending in the locked state^17,27–29^. Within this model, mismatches introduce local energy barriers, while negative supercoiling (favoring DNA unwinding and thus R-loop formation) provides a global negative bias of the energy landscape (Fig. 1b). An important consequence of the model is that even for a fully matching target, by far not every PAM-binding event results in successful target recognition. For example, in absence of a bias, the recognition probability *p*_*rec*_ of a fully matching target of *N* base pairs length would be as low as 1/*N*^28^. The target recognition probability has, however, a direct impact on the required time to search for a target within the vast amounts of non-specific binding sites present in a particular genome. While a target search based solely on three-dimensional (3D) diffusion would prevent extended dwell times at sites far away from the target, the entire genome would, on average, have to be scanned 1/*p*_*rec*_ times for a successful recognition. A purely one-dimensional (1D) diffusion-based target search, on the other hand, would allow rapid repeated revisiting of a target once the enzyme is close to the target, which would compensate for a low recognition probability, as shown for other DNA binding proteins, such as LacI^30^. However, it would at the same time introduce a highly redundant sampling of the whole genome.

Based on these arguments, combined 1D and 3D diffusion, also called facilitated diffusion, should constitute the optimal target search mechanism for CRISPR-Cas surveilance complexes, as it would combine the benefits of both diffusion types. While such a facilitated diffusion mechanism has been proposed for a number of DNA binding proteins^30–32^, partially contradictory results have been obtained for Type I but also other CRISPR-Cas systems. In initial single-molecule DNA curtains experiments, *E. coli* Cascade has been observed to search its target only by 3D diffusion^5^. Binding times to individual PAMs of several seconds were reported. It was proposed that Cascade achieves high search efficiencies simply by discriminating large parts of the DNA due to the mandatory PAM binding. *In-vitro* single-molecule FRET experiments revealed short dwell times (<1s) on PAM-containing DNA. Dwell times increased on DNA containing more PAMs, suggesting a 1D diffusion mechanism for *E. coli* Cascade^33^. A single-molecule *in-vivo* experiment observed considerably lower PAM interaction times of only ∼30ms, with *E. coli* Cascade spending 50% of its search time on DNA and 50% freely diffusing to a new site^34^. While the authors suggested a facilitated diffusion mechanism, 1D diffusion was not directly observed. Lastly, extended 1D diffusion by Cascade from *Thermobifida fusca* over distances of ∼4 kbp was directly observed *in-vitro* with DNA binding times of several seconds^35^. Though these results hint at a possible role of 1D diffusion during target search by Cascade, the relative extent and benefits of the 3D and 1D components during a facilitated diffusion mechanism are not understood. Most importantly, for CRISPR-Cas systems, but also DNA binding enzymes in general, it remains largely unresolved to what extent 1D diffusion is applied to compensate for limited target recognition efficiencies. Furthermore, it has not yet been investigated how target recognition and thus target search are modulated by supercoiling, which is abundantly generated in cellular processes such as transcription and replication^36–38^.

Here, we used a comprehensive set of combined single-molecule fluorescence and magnetic tweezers measurements to uncover the principles of the target search mechanism of St-Cascade. Specifically, we characterized the limited 1D diffusion, the target search efficiency, the time spent between DNA binding and R-loop formation, and the rate at which formed R-loops become locked. Applying different levels of negative supercoiling allowed us to deliberately modulate the target recognition efficiency that could be as low as ∼1% in the absence of supercoiling. However, St-Cascade was found to revisit a target site multiple times per DNA binding event, thereby significantly increasing the absolute search efficiency. We further show that the locking transition is slow compared to R-loop formation steps and directly affects the target recognition in absence of supercoiling. Overall, our study reveals a tight interplay between the target search mechanism, the target recognition probability and the target search efficiency. Importantly, it shows that limited 1D diffusion as well as DNA supercoiling needs to be considered when understanding and modeling (off-)target recognition and target search by CRISPR-Cas surveilance complexes.

## Results

### Investigating DNA binding and R-loop formation by St-Cascade using combined magnetic tweezers and single-molecule fluorescence experiments

To investigate the target recognition process by St-Cascade in detail, we first probed the delay between initial DNA binding and subsequent R-loop formation. We used a microscope setup combining magnetic tweezers and total internal reflection fluorescence (TIRF) microscopy^39–41^. In brief, 2.7 kb dsDNA molecules were attached on one end to the surface of a fluidic cell and on the other end to magnetic beads (Fig. 1c). The DNA molecules were stretched, using the field gradient of a pair of magnets, and the length of the DNA was tracked in real time^42,43^. The DNA molecule contained a single target site situated close to the surface of the fluidic cell. R-loop formation was studied using a previously established magnetic tweezers assay^15^. Rotation of the magnets allowed positive (DNA over-twisting) and negative (DNA untwisting) supercoiling of the DNA molecules, resulting in the reduction of the DNA length due to writhe formation (Fig. 1d)^44–47^. R-loop formation provides an untwisting of DNA by approximately three turns for the full 32 bp R-loop. On negatively supercoiled DNA, the untwisting absorbs a corresponding number of the applied turns such that R-loop formation becomes visible as a sudden length increase of the DNA (Fig. 1e). To observe simultaneously St-Cascade binding and dissociation, the complex was labeled with a single Cy5-dye on the Cas6 subunit (see Methods, Supplementary Figs. 1a, b). Labeling the protein complex did not significantly affect target binding (Supplementary Fig. 1c). To detect DNA-bound complexes, we used TIRF microscopy for which a ∼200 nm deep volume above the sample surface was illuminated with an evanescent field such that only fluorescence of St-Cascade complexes binding near or at the target site was excited. To observe multiple R-loop formation and collapse events on a single DNA molecule, the target sequence for these measurements contained 20 mismatches in the PAM-distal region. Therefore, only unlocked and highly unstable R-loop intermediates of 12 bp length were formed, which spontaneously collapsed after ∼10 s (see DNA length trajectory in Fig. 1f, top right) for the applied negative supercoiling (−6 turns, 0.3 pN). The fluorescence signal from bound St-Cascade was monitored in a fully synchronized fashion^39^. DNA binding was strongly correlated with R-loop formation (fluorescence signal in Fig. 1f, bottom right), i.e., the R-loop formed typically upon binding of a single St-Cascade complex and collapsed upon St-Cascade dissociation. This indicated that R-loop formation occurred very rapidly after St-Cascade binding to the DNA in the vicinity of the target site (within ∼200 nm, corresponding to ∼600 bp). For a minor fraction of R-loop formation events, no Cascade binding was detected alongside R-loop formation due to a limited labeling efficiency of the complex of ∼65% (Supplementary Fig. 2a). Occasionally, Cascade dimers and bleaching were observed as well (Supplementary Figs. 2b, c).

### Cascade forms an R-loop shortly after binding and dissociates shortly after R-loop collapse

We analyzed the time St-Cascade takes from DNA binding to the formation of a 12 bp R-loop, i.e. the search time, as well as the time it takes from R-loop collapse, until its dissociation from the DNA. Since locking of the complex to the target was prevented by the 20 PAM-distal mismatches, binding and dissociation events could be observed repeatedly. Visual inspection of the correlated data suggested dwell times in the sub-second range (Fig. 2a, green and blue arrows, respectively). To quantitatively determine these dwell times, we examined the transition points in the recorded trajectories (from unbound to bound and no R-loop to R-loop states and vice versa) using Hidden Markov modeling^48^ and calculated the respective time differences. The obtained dwell time distributions were rather broad compared to their small positive means (Fig. 2b) due to the limited time resolution in determining the transition points. For the fluorescence measurements, the resolution corresponded to approximately 50 ms due to the camera acquisition rate of 20 Hz. R-loop transitions exhibited even a lower resolution of approximately 140 ms as well as a bias of 13 ms towards larger times, due to the limited relaxation time of the DNA-tethered magnetic bead in aqueous solution (Supplementary Figs. 3, 4; Supplementary Methods 1). Careful consideration of the statistical distributions of the measurement errors and usage of maximum likelihood estimations (Supplementary Methods 2) allowed to extract the mean dwell times (Fig. 2b). On average, St-Cascade was bound to the DNA for *τ*_*R,form*_ = 80 ± 10 ms before forming an R-loop. After R-loop collapse, St-Cascade remained, on average, bound to the DNA for another *τ*_*R,coll*_ = 80 ± 10 ms (Fig. 2c). These results indicated that R-loop formation by St-Cascade occurred rapidly after binding, but not instantaneously.

**Figure 2.**
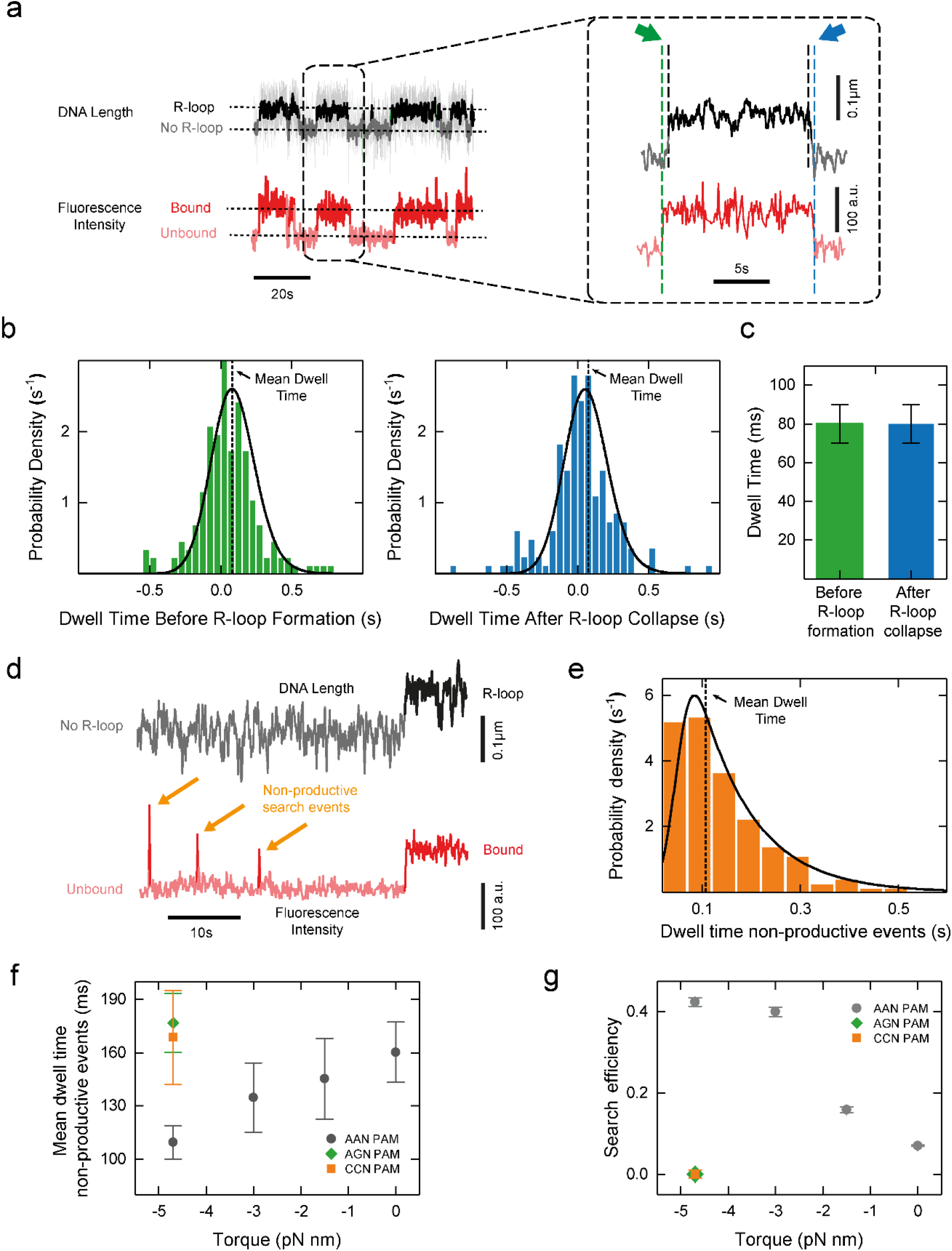
Target search by St-Cascade comprises productive and non-productive search events. **a** Correlated time trajectories of the DNA length and the fluorescence intensity monitoring R-loop formation and St-Cascade binding, respectively. The trajectory on the right is an enlarged view of a single R-loop formation event. Dashed vertical lines indicate the transition points in the trajectories. Colored arrows point at the dwell times before R-loop formation (green) and after R-loop collapse (blue). The trajectory was taken at 120 Hz and smoothed to 3 Hz. The fluorescence trajectory was recorded at 10 Hz. **b** Histograms of the measured dwell times between St-Cascade binding and R-loop formation (left, *N* = 173) as well as R-loop collapse and St-Cascade dissociation (right, *N* = 164). Black solid lines represent the maximum likelihood estimates of the data and dashed lines indicate the mean dwell times. **c** Mean dwell times for R-loop formation and after R-loop collapse from maximum likelihood estimation shown in b. **d** Correlated time trajectories of the DNA length and of fluorescence signal revealing short non-productive search events in which St-Cascade binds to the DNA and dissociates without R-loop formation, reflected by the absence of a length change in the DNA length trajectory. The DNA length was recorded at 120 Hz and smoothed to 3 Hz, while the fluorescence trajectory was taken at 10 Hz. **e** Dwell time histogram of the non-productive search events for г=-4.7 pN nm (F=0.2 pN, -6 turns). The black line represents the maximum likelihood estimate of the data (*N* = 142). **f** Mean dwell times of non-productive search events as function of the applied torque for targets with the cognate (AAN) PAM and two non-cognate (AGN and CCN) PAMs. **g** Search efficiency, i.e., the percentage of R-loop forming events within all binding events, as function of torque for targets with the indicated PAMs.

### Cascade exhibits non-productive search events with torque-dependent dwell times

Within the observed dwells between DNA binding and R-loop formation, it is unclear whether St-Cascade rests on the target PAM before forming an R-loop or actively searches for the target PAM among the multitude of neighboring PAMs. To discriminate between the two possibilities, we inspected our fluorescence trajectories more closely. In addition to long-lived Cascade binding events that corresponded to formed R-loops, we also observed many short-lived, non-productive binding/search events for which no R-loops were formed (Fig. 2d). To better understand the fate of St-Cascade within the dwells, we also determined the dwell times, *τ*_*no R−looP*_, of these events. The dwell times were only slightly larger than the time resolution of the fluorescence measurements and we again used maximum likelihood estimation to obtain mean dwell times from the measured distributions (Fig. 2e). This analysis could account for very short events that remained undetected, as confirmed by analyzing simulated trajectories (Supplementary Fig. 5; Supplementary Methods 3). We repeated the analysis at various supercoiling levels (Supplementary Fig. 6), represented by the mechanical torque, which could be set by changing the applied stretching force (Methods). At low supercoiling levels (close to zero turns), R-loop formation is not visible as a length change, because of lacking writhe formation (see Fig 1d). Using a fully matching target, DNA binding events could be seen in the fluorescence trajectories as long-lived binding events following locking of the R-loop. They were, however, clearly distinguishable from the short-lived non-productive search events (Supplementary Fig. 7). Interestingly, the dwell times of the non-productive search events strongly decreased with increasing negative supercoiling from 160 ms to 110 ms (Fig. 2f). Assuming that non-productive search events correspond to non-specific DNA binding and that the dissociation rate *K*_*diss*_ for non-specifically bound St-Cascade is independent of supercoiling, the most probable explanation for the decreasing dwells of non-productive search events is that competing productive R-loop formation becomes favored at negative supercoiling as previously observed^15,16,49^. Therefore, the observed torque-dependent dwells hint at a 1D search process for which any observed St-Cascade molecule bound to the DNA can either non-productively dissociate or form an R-loop. To corroborate that the source of the decreasing dwell times was indeed facilitated R-loop formation rather than a supercoil-dependence of non-specific binding, we repeated the measurements with two target variants containing either an AGN or a CCN PAM, which reduced or fully inhibited R-loop formation (Supplementary Fig. 8). At high negative supercoiling, the dwell times of non-specific St-Cascade binding were, for both targets, not reduced but instead corresponded to the dwell times measured for the cognate PAM (AAN) at zero supercoiling, where R-loop formation is also inhibited (Fig. 2f). Thus, we concluded that the reduced dwell times were indeed caused by the competing R-loop formation process rather than a torque-dependence of non-specific binding.

### St-Cascade exhibits a torque-dependent search efficiency

To obtain direct support for the hypothesis that negative supercoiling favors R-loop formation at the expense of non-productive dissociation, we directly determined the efficiency of the search process *E*_*search*_ from the number of productive (R-loop formation) and non-productive (dissociation without R-loop formation) search events. As expected, increasing negative torque increased the efficiency of the search process. While in absence of supercoiling, successful R-loop formation was observed for only 7±1% of all binding events, it was seen for more than 40% of the events at high supercoiling levels (Fig. 2g). Given that the illuminated DNA segment of ∼200 nm length contained ∼70 PAMs, a target search based on pure 3D diffusion that selects a single PAM would have much lower search efficiencies than observed in our experiments. This further supports that St-Cascade searches for the target by scanning the permissive PAMs using 1D diffusion when bound to the DNA.

### St-Cascade uses limited 1D diffusion for its target search

To directly verify that 1D diffusion is part of the target search mechanism, we followed the movement of St-Cascade on a 15 kbp DNA fragment, which did not contain a target site. For that, we employed a cylindrical magnet for lateral DNA stretching. At sufficient distance from its symmetry axis, it produced a field gradient with a strong lateral component thus conveniently enabling lateral pulling of the magnetic bead (Fig. 3a). Using TIRF microscopy, we monitored the interaction of the Cy5-labeled St-Cascade with the non-specific DNA. As predicted, we observed short-lived binding events with visible diffusion along the DNA, spanning several hundred nanometers (Fig. 3b). After tracking the movement of St-Cascade for a large number of events, we determined the mean-squared displacement ⟨*x*^2^⟩ as a function of time, which increased linearly, in agreement with 1D diffusion along DNA, for which ⟨*x*^2^⟩ = 2*Dt*. A fit of the data provided a 1D diffusion coefficient *D* = 0.014 ± 0.004 µm^2^/s (corresponding to 0.1 kbp^2^/s) (Fig. 3c). Furthermore, as a control, we determined the dwell times of all individual binding events, yielding an average of *τ*_*no R−looP*_ =170 ± 10 ms (Fig. 3d). This equaled, within errors, the dwell times obtained previously for the non-specific targets (see Fig. 2f). From these parameters, we calculated that St-Cascade scans, on average, a distance of 90 ± 10 nm (270 ± 30 bp) per binding event, containing approximately 35 AAN PAMs (with one AAN PAM every ∼8 bp). Approximating the 1D diffusion as a random-walk from one AAN PAM to the next with negligible time spent in between as suggested before^33^, we obtained that St-Cascade, on average, performs 320 ± 90 steps per binding event and stays bound to a single PAM for *τ*_*PAM*_ = 0.5 ± 0.2 ms. These data provide direct support for a facilitated diffusion model in which St-Cascade binds DNA by a 3D diffusion mechanism and then scans a limited region in 1D.

**Figure 3.**
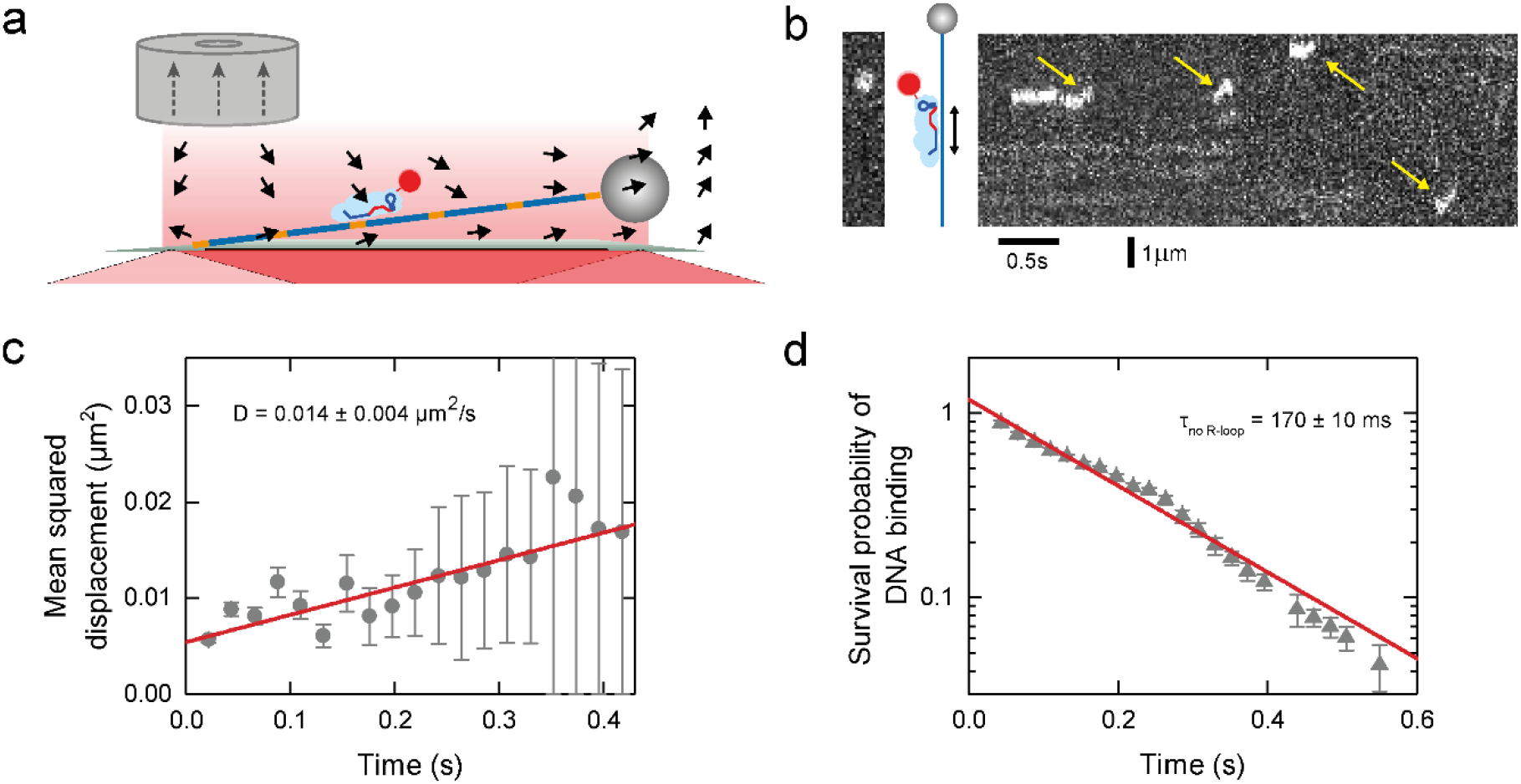
St-Cascade scans DNA by combining 3D and 1D diffusion. **a** Scheme of lateral DNA stretching with magnetic tweezers and simultaneous TIRF microscopy. Orange sections on the DNA represent PAMs. Black arrows represent the magnetic force due to magnetic field gradient. Grey dotted arrows depict the magnetization of the magnet. **b** Kymograph (right) of the fluorescence intensity along the DNA revealing short-lived binding events and limited 1D diffusion along DNA (yellow arrows). The far left image depicts a single snapshot of the measurement with St-Cascade visible as a white dot. **c** Mean squared displacement as function of time obtained from multiple binding events (blue circles, *N*= 116). A linear fit to the data (red line) provided a diffusion coefficient of *D* = 0.014 ± 0.004 µm^2^/s (see main text). **d** Survival probability of the DNA binding events as function of time (*N*=116). Fitting an exponential decay to the data (red line) provided a mean binding time of 170 ± 10 ms.

### St-Cascade scans matching targets multiple times to increase the search efficiency

The established 1D diffusion by St-Cascade suggests that the complex revisits a target site multiple times before successful recognition. The measured search efficiency per DNA binding event should thus be considerably larger than the actual recognition probability of the matching target per single target site encounter. To estimate the target recognition probability and integrate the different obtained data sets, we established a kinetic model for the target search of St-Cascade (Supplementary Methods 4): we described the target search as a random walk on a 1D lattice containing *N*_*PAM*_ lattice points, representing cognate AAN PAMs (Fig. 4a; Supplementary Fig 9a). One of the PAMs was selected to contain the matching target. After initial random binding to a PAM on the lattice, the complex can successively either take a step to one of its two neighboring PAMs with a stepping rate *K*_*step*_ = 1/*τ*_*PAM*_ = 1.9 ms^-1^ or dissociate with the measured dissociation rate *K*_*diss*_ = 1/*τ*_*non−spec*_ = 5.7 s^-1^ from non-specific DNA.

**Figure 4.**
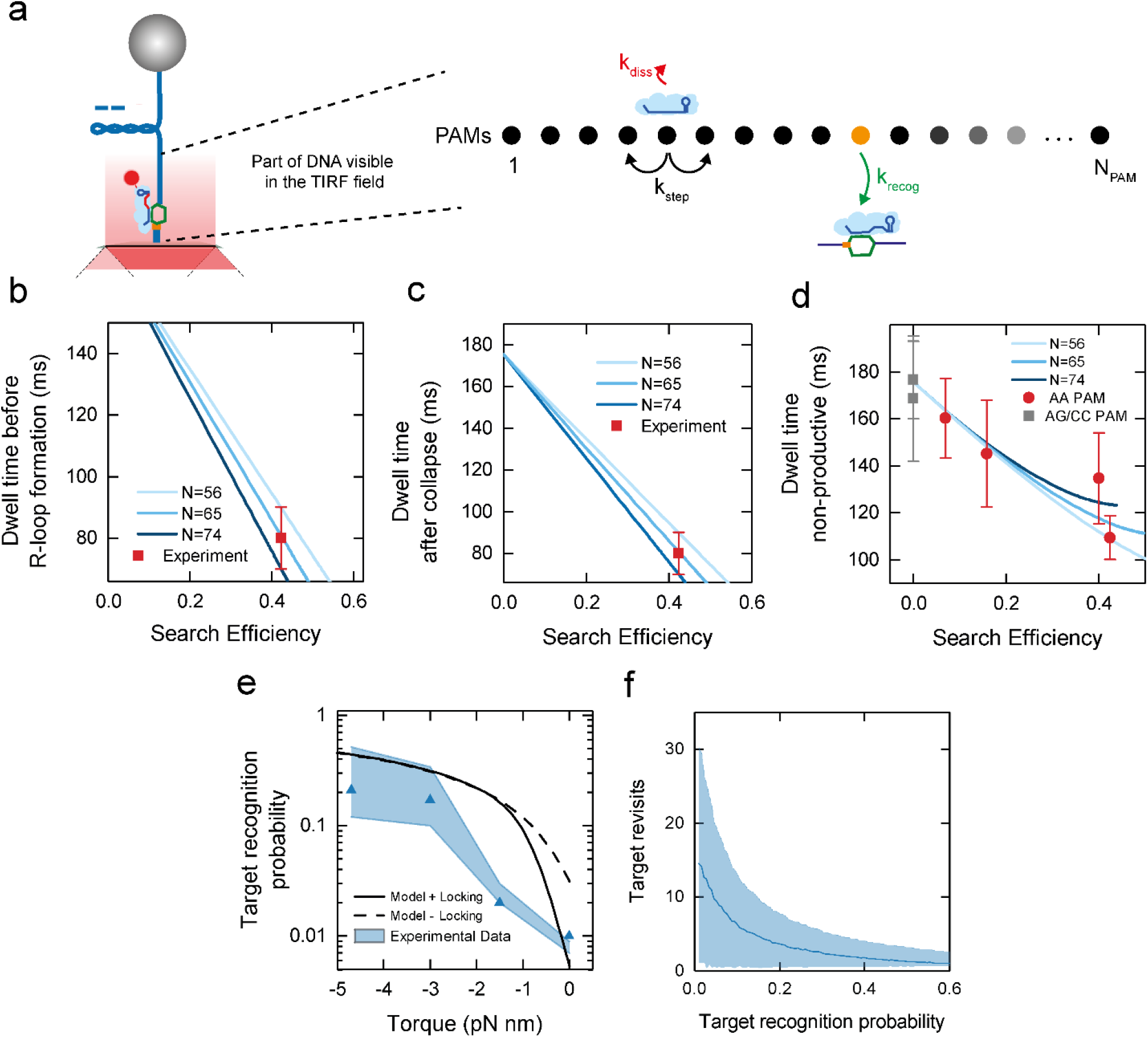
St-Cascade target search modeled as a random walk on 1D lattice containing *N*_*PAM*_ PAMs. **a** Schematic representation of the target search model. **b** Dwell time *τ*_*R,form*_ on DNA before R-loop formation as function of the obtained search efficiency for varying lattice sizes (solid lines). The respective experimental value (red square) agrees with the model for a lattice size of *N*_*PAM*_ = 65±9. **c** Dwell time *τ*_*R,coll*_ on DNA after R-loop collapse as function of the obtained search efficiency for varying lattice sizes (solid lines). The experimental value (red square) agrees with the model for a lattice size of *N*_*PAM*_=65±9. **d** Dwell times of non-productive search events modelled for varying lattice sizes as function of the search efficiency (solid lines). For comparison, the experimental values for a cognate PAM (AAN, red circles) and two non-cognate PAMs (AGN/CCN, grey squares) are shown. **e** Estimated target recognition probability *p*_*recog*_ (per target encounter) as function of torque. Results are shown for a lattice size of *N*_*PAM*_ = 65 (blue triangles) where the shaded blue area represents the uncertainty given the error of *N*_*PAM*_ = 65±9. **f** Average number of target site revisits during a productive target search event as function of *p*_*recog*_ (blue line). Site revisits were obtained from random walk simulations with 10,000 search events for each data point. Searches started at the target PAM. The blue shaded area depicts the standard deviation of the resulting distributions of target revisits.

Upon visiting the PAM with the matching target, St-Cascade can recognize this target with rate *K*_*recog*_. The recognition probility per target binding event is then given as *p*_*recog*_ = *K*_*recog*_/(*K*_*recog*_ + *K*_*diss*_ + *2K*_*step*_). We derived a solution to this model and calculated the kinetics of the formation of productive and non-productive search events that were terminated by successful target recognition or Cascade dissociation, respectively (Supplementary Figs. 9b, c). This allowed us to extract the corresponding dwell times *τ*_*no R−loop*_, *τ*_*R,form*_ and *τ*_*R,coll*_ as well as the target search efficiency *E*_*search*_ (Supplementary Methods 4). Unknown parameters of the model were the recognition probability, *p*_*recog*_, and the lattice size *N*_*PAM*_, the latter being determined approximately by the length of the illuminated DNA region (Fig. 4a). To determine the lattice size more accurately, we carried out model calculations of productive events for a range of different values of both parameters and compared the obtained dwell times to the experimental results of *τ*_*R,form*_ and *τ*_*R,coll*_. They were consistently described by lattice lengths between 56 and 74 PAMs, given the experimental errors (Figs. 4b, c). This agreed with the experimentally adjusted depth of the evanescent field of 150 to 200 nm, given a cognate PAM every ∼8 bp. We repeated the model calculations for non-productive search events and found that the dependence of the dwell times, *τ*_*no R−loop*_, on the search efficiencies was accurately described by our model for the selected lattice lengths (Fig. 4d). This suggested that our model can quantitatively describe the target search process by St-Cascade. Finally, we used the model and the selected lattice lengths to calculate estimates of the target recognition probability as function of torque (blue triangles and shaded area, Fig. 4e).

The recognition probability, *p*_*recog*_, ranged from ∼1% in the absence of supercoiling to ∼25% at the highest negative supercoiling. It was thus always significantly below 1% but also significantly lower than the measured search efficiencies (see Fig. 2g). Notably, for the low target recognition probabilities, site revisits due to the 1D diffusion process rescued the search efficiencies considerably (from 1% (target recognition) to 7% (search efficiency) at zero torque). The number of site revisits depended on the target recognition probability, i.e., the applied supercoiling level (Fig. 4f), ranging from ∼15 revisits in absence of supercoiling (*p*_*recog*_ = 1%) to ∼3 revisits at the highest negative supercoiling (*p*_*recog*_ = 25%).

Above, we determined the target recognition probability by modeling the target search efficiency. However, it can also be directly predicted using the target recognition model based on the discrete 1D energy landscape described in the introduction^16,17,27,29^ (Fig. 1b; Supplementary Fig. 10). Agreement of both models would provide a direct link between the target recognition and target search mechanisms. Assuming that full R-loop formation instantaneously promotes locking, the target recognition probability *p*_*recog*_ is equivalent to the formation probability of a full R-loop after arrival at the correct PAM, *p*_*Rloop*_. We calculated theoretical values for *p*_*Rloop*_ as function of torque and R-loop length using a constant torque-induced free energy bias per base pair (Supplementary Figs. 11, 12; Supplementary Methods 5) and compared it to *p*_*recog*_ obtained from the experimental data (Fig. 4e, black dotted line). Within error, agreement was obtained for elevated negative supercoiling. However, *p*_*Rloop*_ considerably overestimated *p*_*recog*_ at lower negative supercoiling.

### The R-loop locking transition limits the target recognition in absence of supercoiling

The observed discrepancies between *p*_*recog*_ and *p*_*Rloop*_ from the target recognition model at low negative supercoiling may be due to a rate limiting locking transition. If locking of full R-loops is sufficiently slow, they can still collapse, particularly at low negative supercoiling at which they are little stabilized. Slow locking would thus lead to a reduced target recognition probability. To test this idea, we sought to directly determine the intramolecular locking rate *K*_*lock*_ of a fully formed R-loop by St-Cascade. To this end, we monitored transient, short-lived R-loop states with magnetic tweezers, and used them as probes for the locking transition. Particularly, we applied a target with one internal mismatch at position 17 (Fig. 5a). Since R-loop locking is very fast for a target with a fully matching PAM-distal end, we slowed down this process by introducing 1, 2, or 4 additional mismatches in this region. On these targets, three different R-loop states were observed: an unbound state, an intermediate state with a 16 bp-long R-loop, and an (almost) full R-loop (Fig. 5a). As long as the R-loop remained unlocked, we observed a rapid sampling between these different states. Upon locking, the sampling was suddenly stopped and the R-loop remained stable in the full state (orange section in Fig. 5a). In the unlocked full R-loop state, R-loop collapse towards the intermediate state competes with R-loop locking. The apparent locking rate in the full R-loop state is then given by the rate, *K*_*coll*_, at which unlocked R-loops collapse and the probability, *p*_*lock*_, that a full R-loop becomes locked once formed (Supplementary Methods 6):

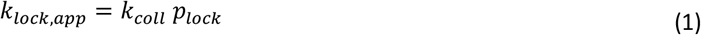

**Figure 5.**
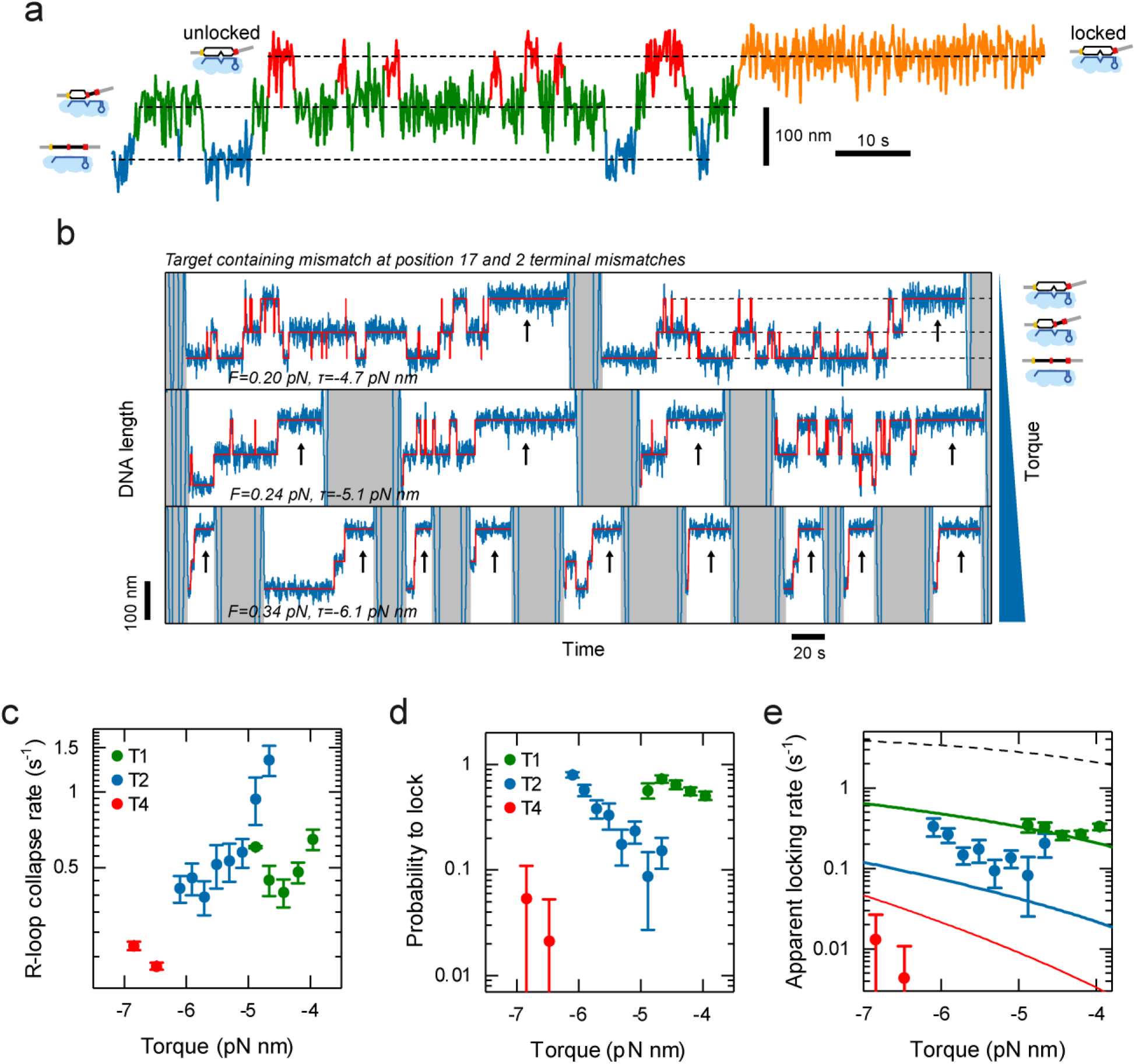
Investigating the target recognition process. **a** Characteristic time trajectory of transient R-loop states on a target with a mismatch at position 17 and varying numbers of PAM-distal mismatches. Visible are the unbound state (blue), the intermediate R-loop state (green), and the full R-loop states (red and orange). In the full state, the R-loop can either be unlocked such that it can collapse back to the intermediate state (red) or be stably locked (orange). b Characteristic time trajectories (blue) for a target containing a mismatch at position 17 and two PAM-distal mismatches recorded at different negative supercoiling. Idealized trajectories from Hidden Markov modeling are shown in red. The trajectories show repeated cycles of sampling the different R-loops states until locking occurs (black arrows). After each locking event, R-loop collapse and dissociation of St-Cascade is enforced by applying high positive torque (grey areas). With increasing torque, the R-loop becomes more rapidly locked. c,d The rate *K*_*coll*_ at which full unlocked R-loops collapse back to the intermediate state and the probability, *p*_*lock*_, that an R-loop in the full state becomes locked measured for 1,2 and 4 of terminal mismatches (T1, T2, T4) as function of torque. e The apparent locking rate, at which an R-loop in the full state becomes locked as function of torque calculated from *K*_*coll*_ and *p*_*lock*_ for the 3 different targets. The experimental data is depicted by filled circles while solid lines represent a global fit to the data according to a simple locking model (Eq. 1). The model prediction for a target without terminal mismatches is shown as a black dashed line. All trajectories were taken at 120 Hz and smoothed to 3 Hz

Both parameters could easily be inferred from trajectories that we recorded for targets with one, two, or four terminal mismatches (T1, T2, T4). We repeated these measurements at varying torques to determine a potential torque dependence (Fig. 5b). The locking rate, which strongly differed for the different targets, was limiting the applicable torque range. Generally, the R-loop collapse rates decreased with increasing negative torque, since the DNA untwisting stabilized the full R-loop state (Fig. 5c, Supplementary Fig. 13). The probability, *p*_*lock*_, that a full R-loop became locked rather than collapsed, decreased strongly as more terminal mismatches were introduced (Fig. 5d) and increased with increasing negative torque. When finally calculating *K*_*lock,app*_, we obtained an increase of the apparent locking rate with increasing negative torque (Fig. 5e), similar to *p*_*lock*_. We also observed a strong decrease of the apparent locking rate with an increasing number of mismatches, as seen for *p*_*lock*_. The strong dependence of *K*_*lock,app*_ on the mismatch number suggests that the R-loop has to extend over its full length of 32 base pairs in order to efficiently promote locking. However, the unlocked R-loop does not always extend to the last available base pair, but dynamically samples also shorter lengths as determined by the energy landscape (see Fig. 1b). This sampling is changed by the bias of the energy landscape. Increasing supercoiling, for example, favors longer R-loops. Thus, locking should become favored with increasing negative supercoiling, in agreement with the observed behavior for *K*_*lock,app*_. To test whether a supercoiling-dependent stabilization of extended R-loops dominates the observed torque dependence, we modelled this process by assuming that locking can only occur for a fully extended 32bp R-loop. The rate *K*_*lock,app*_ is then given by the probability, *p*_32_, that the R-loop extends to its maximum length of 32 bp, multiplied by the ‘true’ rate, *K*_*lock*_, at which locking occurs, when the R-loop is fully extended:

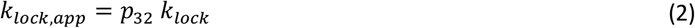

The probability *p*_32_ can easily be approximated using the simplified torque-dependent energy landscape into which penalties for the terminal mismatches were incorporated (Supplementary Fig. 14; Supplementary Methods 7). Fitting the resulting prediction to the experimental data allowed to approximately describe the observed torque-dependence of *K*_*lock,app*_ (Fig. 5e, solid lines). The fit provided a locking rate of *K*_*lock*_ =6 ± 2 s^-1^, i.e. a locking transition time in the range of 100 ms, as well as a mean mismatch penalty of 2.7 ± 0.3 k_B_T for terminal mismatches, which agreed with the magnitude of the base-pairing energies of DNA base pairs. Furthermore, the model could describe the observed dependence on the mismatch number. Only for the four terminal mismatches (T4), larger deviations were observed, possibly due to local variations of the mismatch penalties. For the absence of PAM-distal mismatches, the model predicts apparent locking rates >1 s^-1^ (Fig. 5e, dashed line), which would be difficult to characterize, given the obtained values for *K*_*coll*_.

Overall, the description of the torque and mismatch dependence of *K*_*lock,app*_ supports the idea that R-loop locking only occurs for a fully extended R-loop at the intramolecular locking rate *K*_*lock*_. Incorporating an additional locking step into the random-walk model for target recognition and assuming a base pair stepping rate *K*_*bp*_ (see Fig. 1a; Supplementary Methods 7) in the millisecond range^17^, we could obtain a prediction for the target recognition probability *p*_*recog*_ as function of torque that included locking (Fig. 4e, black line). While it did not deviate from *p*_*Rloop*_ at higher negative torque, it better described the strongly reduced recognition probability in absence of torque.

## Discussion

In this study we used a comprehensive set of single-molecule experiments in order to dissect the target search mechanism of the St-Cascade surveillance complex. Particularly, we applied single-molecule fluorescence microscopy to follow the 1D diffusion of single complexes along stretched DNA, magnetic tweezers to monitor R-loop formation, collapse and locking as well as correlated measurements of the two techniques to follow the dwells between DNA binding and R-loop formation and to determine the search efficiency. Kinetic modelling was applied to integrate the different data sets. Notably, the possibility to apply different supercoiling levels was a helpful for modulating the target recognition probability upon target binding by Cascade and monitor its impact on the measured search efficiencies and dwell times.

From these results, a detailed picture of the target search process emerged (Fig. 6): Cascade uses 3D diffusion to bind non-specifically to DNA for 100-200 ms. There, it undergoes limited 1D diffusion along the DNA over a mean length of 270 basepairs. During the 1D search on our DNA construct, it rapidly interrogates ∼35 PAMs within this region employing very short PAM binding times of only 0.5 ms per PAM. If a target site is nearby, it is repeatedly revisited during diffusion until it is recognized or the complex dissociates from the DNA to continue its 3D search.

**Figure 6.**
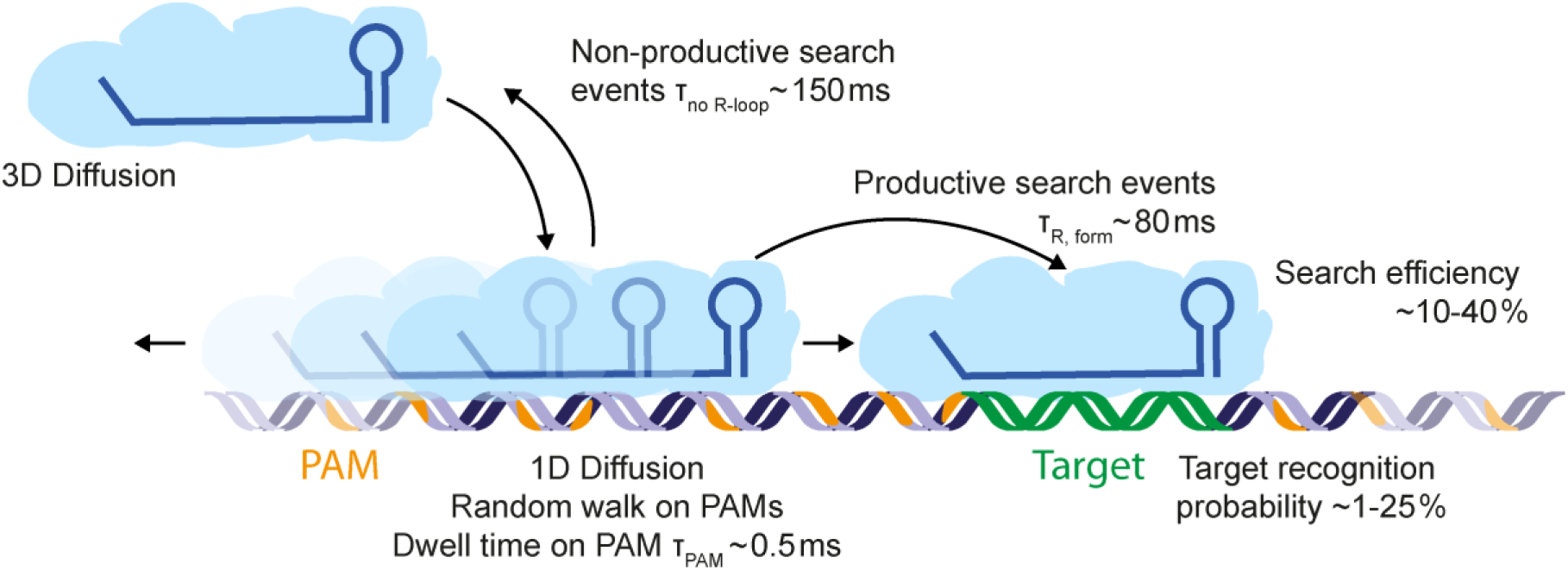
Schematic facilitated diffusion search process by St-Cascade. Following 3D diffusion, St-Cascade binds to DNA and scans PAMs in a 1D random walk process. Each PAM is interrogated for approximately 0.5ms. Productive 1D search events that end in the formation of a full R-loop take on average 80 ms. Non-productive search events end after approximately 150 ms by dissociating from the DNA and proceeding to 3D diffusion. The duration of non-productive search events depends on the torque applied on the DNA where adding negative torque reduces the duration of the search due to facilitated R-loop formation. In case of encountering the correct PAM, R-loop formation, and thus target recognition, occurs in less than 25% of the cases. However, St-Cascade revisits the PAM multiple times, leading to a significant increase in overall search efficiency up to 40% at negative torque.

The target recognition probabilities per single target encounter are rather low ranging from <1% to ∼25% at zero and high negative supercoiling, respectively. Probabilities well below 100% are in agreement with the required reversibility of the R-loop formation process which forms the basis of the rapid scanning of the available sequence space. Site rescanning during 1D diffusion rescues, however, the search efficiencies to values between 7% and 42%. The observed target search behavior is thus a compromise between a pure 3D search mechanism that allows to quickly sample all distant regions of an entire genome and a 1D search mechanism that very thoroughly probes the existence of a site within a given region and avoids void time off the DNA. It thus appears that the limited 1D diffusion during the observed facilitated target search is used to compensate the low target recognition efficiencies. This suggests that the combination of 3D and 1D diffusion is not only applied to reduce the time for 3D diffusion off the DNA but rather to make the local search more efficient. This together reduces the overall search time. Similarly to St-Cascade, the bacterial DNA binding protein LacI has very recently been shown to have target recognition probabilities that were significantly lower than 100%. This was due to hops between DNA grooves which interrupt the DNA sliding and scanning along the helical path^30^. Also here, a limited 1D search on top of the 3D pathway should help to rescue the final search efficiency making it to a much more common mechanism among DNA binding enzymes.

Results of experimental and theoretical studies on classical DNA binding enzymes, such as Type II restriction endonucleases, RNA polymerases and transcription factors, have indicated a 1D diffusion length of 10-100 bp for an optimal search of specific targets^50–53^. For St-Cascade we observed here a mean distance of ∼270 bp which is considerably larger than the proposed optimum. We attribute this to the almost exclusive limitation of the search process to cognate PAM sequences^33^. This enables St-Cascade to scan, within a given region, much fewer potential targets compared to classical DNA binders that would probe the full sequence space. For a limited sequence space, the optimum for a 1D search would likely be shifted to larger distances as seen for Casacade. Specific DNA recognition by classical DNA binders occurs more as an allosteric single-step process^54^ and therefore much more rapidly than the slow multi-step, random walk-like target recognition by CRISPR-Cas effectors. Reducing the target-search for the latter mainly to PAM sequences, at which the adjacent target gets thoroughly probed, will nonetheless ensure a rather efficient and rapid target search. Accelerated target search represents thus an additional benefit for the usage of PAM elements beyond the discrimination of self vs. non-self. This should be taken into account when engineering effector variants with shorter or no PAM sequences. For the latter, the probed sequence space will be increased by a factor of ∼4^*n*^, with *n* being the number of unique base pairs of the PAM, potentially providing a tremendous increase for the search times of these variants.

We further showed that the target recognition efficiency and thus the search efficiency are strongly modulated by DNA supercoiling. Particularly, the recognition was rather inefficient in absence of supercoiling. This behavior could be reproduced by modelling the target recognition as a random walk of the forming R-loop in which the applied negative supercoiling biases the energy landscape of R-loop formation and increases the probability of R-loop formation. This provides further evidence for the established target recognition model for CRISPR-Cas effector complexes and strand exchange reactions in general^16,17,27–29^. Most importantly, it demonstrates that target search and the target recognition mechanism are tightly linked. The target recognition probability in absence of supercoiling was additionally reduced by the transition that locks the full R-loop irreversibly in a stable state before DNA cleavage. The locking transition was found to be ∼150-fold slower than the R-loop stepping rate, such that it provided a significant effect.

The strongly reduced target recognition efficiency in absence of supercoiling suggests that CRISPR-Cas systems have evolutionary been optimized to support efficient recognition of invaders with supercoiled DNA. This is supported by the discovery of a novel anti-CRISPR protein, which nicks plasmid DNA to release supercoils in order to strongly reduce the targeting efficiency of CRISPR-Cas9^55^ as well as the torque-dependent post-cleavage behavior of Cas9 itself^56^. The genomes of eukaryotic cells are however considerably less supercoiled than prokaryotic cells. For genome engineering applications in eukaryotes, one should therefore bear in mind that the used CRISPR-Cas effectors are typically not optimized for the corresponding supercoiling conditions at least regarding the target search efficiency. Targeting is thus expected to occur significantly slower. We note, however, that the reduced bias on R-loop formation in absence of supercoiling can increase the targeting specificity since the length of the seed region for R-loop formation by Cascade was shown to be mainly determined by supercoiling^16^. We therefore believe that for each applied effector complex it would be necessary to determine the optimum bias that provides high specificity and little compromised search efficiency. This would allow to use low concentrations of effector complexes ensuring minimal side effects. The suggested optimization could be for example established using rational engineering of effector variants^57–67^ and corresponding characterization *in vitro* and *in vivo*.

Overall our findings suggest that DNA supercoiling as well as limited 1D diffusion need to be considered when understanding and modeling (off-)target recognition and target search by CRISPR-Cas enzymes. Moreover, they may contribute to a better understanding of the search mechanisms employed by other sequence-specific targeting proteins employing a combination of 1D and 3D pathways.

## Materials and Methods

### DNA substrates

Double-stranded DNA constructs for magnetic tweezers experiments with lengths of 2100 bp and 2700 bp were prepared as previously described^16^. A single copy of a given Cascade target including a AAN PAM was cloned into the SmaI site of a pUC19 plasmid. From the plasmid, a 2.1 kbp or 2.7 kbp fragment, including the St-Cascade target site (see Supplementary Table 1) was amplified by PCR, using primers in which either a NotI or a SpeI restriction enzyme site was introduced. After digestion with NotI and SpeI, the fragment was ligated at either end to ∼600-bp-PCR fragments containing multiple biotin (SpeI site) or digoxigenin (NotI site) modifications^68^ to allow the tethering to coated magnetic beads and the coated flow cell of the magnetic tweezers setup. Ligated DNA constructs were separated with agarose gels from other reaction products and purified by gel extraction (NucleoSpin, Macherey-Nagel) by avoiding any exposure to ethidium bromide or UV light.

For diffusion measurements, a 17 kbp double-stranded DNA construct was prepared from three individual pieces. A 9.5 kbp fragment of a pUC18 variant (pUC18-48×601-197)^69^, a 4.3 kbp PCR fragment of plasmid pSC60b^70^ and a 3.2 kbp fragment of Lambda DNA were ligated. Attachmend handles were used as described above.

### Cloning of Cascade-Cas6-V76C and Cas8e

#### Streptococcus thermophilus

DGCC7710 Cascade complex naturally encodes two cysteine residues in Cas8e protein (previously known as Cse1 or CasA) (C252 and C262). To avoid labeling of C252 and C262 in Cas8e, double mutation C525S-C262S was introduced in Cas8e performing PCR from pCDF-Duet vector encoding Cascade effector (with oligonucleotides IR76 and IR77, Supplementary Table 1). Next, cysteine mutation was introduced to V76C residue of Cas6 in the same pCDF-Duet vector (with oligonucleotides IR92 and IR93, Supplementary Table 1). Cas8e was cloned as a standalone gene to pBAD24 vector using NcoI and XhoI cleavage sites fusing Cas8e with C-terminal His_6_-tag (Supplementary Table 1). All primers ordered as desalted from Metabion. All constructs were confirmed by Sanger sequencing.

### Expression and purification of proteins

#### Streptococcus thermophilus

DGCC7710 Cascade complex was heterologously expressed in *E. coli* BL21 (DE3) cells using pACYC pCRh encoding homogeneous CRISPR region ^20^, pCDF-Duet vector with triple mutant Cascade effector Cas8e-C252S-C262S-Cas6-V76C and pBAD-Cas7-C-His (Supplementary Table 1). Upon expression, LB broth (Formedium) supplemented with ampicillin (25 mg/ml), chloramphenicol (17 mg/ml), and streptomycin (25 mg/ ml). Cells were grown at 37°C, 200 rpm to OD_600nm_ of 0.5-0.7 and expression was induced with 0.2% (w/v) L(+)-arabinose and 1 mM IPTG for 3 h. The Cascade complex was purified by Ni^2+^-charged HiTrap column (GE Healthcare) followed by Superdex 200 (HiLoad 16/60; GE Healthcare) as described before^4^, but heparin column was exchanged to Q Sepharose Fast Flow (GE Healthcare) (starting buffer: 20 mM Tris-HCl pH=8.0, 100 mM NaCl, elution buffer: 20 mM Tris-HCl pH=8.0, 1000 mM NaCl). After the last step, it was observed that Cas8e C252S-C262S mutant protein is not present in the complex (Supplementary Figure 15a), therefore wt Cas8e was purified separately expressing it from pBAD-Cas8e-C-His vector (Supplementary Table 1). Upon expression of Cas8e, LB broth (Formedium) supplemented with ampicillin (50 mg/ml) and the cells were grown at 37°C, 200 rpm to OD_600nm_ of 0.5-0.7 and expression was induced with 0.2% (w/v) arabinose. The Cas8e was purified by Ni^2+^-charged HiTrap column (GE Healthcare) followed by heparin column (GE Healthcare) following the same protocol as for Cascade complex, described before^4^. Wt Cas8e-C-His was supplemented to the mutated Cascade-Cas6-V76C complex lacking Cas8e and the cleavage reaction was performed with pSP1-AA (with a protospacer) and pSP3-AA (without a protospacer) supercoiled plasmids in a presence of Cas3 nuclease-helicase and other required components (Supplementary Figure 15b)^4^. Cas3 protein for the cleavage assays was purified as described previously^20^, but the heparin column was not used.

### Fluorescent labeling of Cascade

A Cy5 fluorophore with a malemeide linker was attached to the cysteine within the Cas6 protein of the Cascade effector complex, which had a V76C mutation. First, the storage buffer (20mM Tris-HCl pH 8.0, 500mM NaCl, 50% Glycerol) was exchanged with start buffer (20mM NaP pH 7.0 + 50mM NaCl) in a 100k size exclusion column (Amicon). Cy5-malemeide (Lumiprobe) was dissolved in DMSO. The protein (5µm) was incubated at room temperature in the dark for three hours with a 20x molar excess of Cy5. The labeled protein was then purified from excess fluorophores via a 100k size exclusion column using elution buffer (20mM NaP pH 7.0 + 1M NaCl) until the remaining dye concentration of the flow-through was negligible, as determined with a Nano-photometer (Implen). Finally, the labeling efficiency was measured to be approximately 65% using said Nano-Photometer. Before storage at -20°C, the elution buffer was exchanged with storage buffer.

### Functionalization of the flow cell and the magnetic beads for combined magnetic tweezers and TIRF fluorescence experiments

For binding the DNA substrates to the microfluidic flow cells, glass slides were passivated and functionalized with biotin.

To this end, glass slides (Menzel) were first cleaned by sonication in acetone. Following a further sonication step in KOH (5M), the slides were rinsed with deionized water and MeOH before drying with N_2_. For passivation, the slides were first incubated in 150ml MeOH, 7.5ml acetic acid and 1.5ml aminopropylsilane. The glass slides were then coated with a mixture of mPEG (Rapp Polymere) and biotinylated mPEG (10:1) dissolved in sodium bicarbonate at pH 8.5 and incubated overnight. The slides were stored under vacuum conditions at -20°C.

For tethering the DNA to superparamagnetic beads, beads with a 0,5µm diameter with a carboxylic acid activated surface (Ademtech) were coated with anti-digoxigenin. First, the beads (1mM) were activated by resuspending in MES (25mM) and then incubating with EDC (0.5mg/ml in MES) at 40°C for 10’ while shaking. Anti-digoxigenin (50µg for each mg of beads) was added to the solution and shaken for 2h at 40°C. For passivation, BSA (0.5mg/ml) was added and the solution was incubated in a shaker at 40°C for another 30 minutes. Lastly, the beads were washed with Ademtech storage buffer in a magnetic rack and stored at 4°C.

### Combined magnetic tweezers and TIRF microscopy experiments

The single-molecule measurements were performed in a custom-built magnetic tweezers setup^43^ with integrated TIRF microscopy^39^.The simultaneous DNA length measurement and the recording of fluorescent images was carried out in fully synchronized manner. Prior to the measurements, DNA constructs were bound at their biotinylated end to magnetic beads of 0.5µm diameter (Ademtech), which provided a reduced background in the TIRF measurements due to their small size by minimizing backscattering of the excitation light. Subsequently the bead-tethered DNA molecules were flushed into the fluidic cell of the setup allowing the anchoring of the biotin-modified end to the streptavidin-coated surface of the cell. After removing unbound beads by flushing, force was applied by lowering the magnets towards the flow cell and suitable DNA tethered beads of the expected length were selected. For magnetic tweezers measurements, the sample was epi-illumination was provided by infrared light using a laser ignited XE-plasma lamp (EQ-99-FC, Energetiq) and near-infrared (>770nm) long-pass filter. The DNA length was determined from the axial position of a selected DNA tethered magnetic bead with respect to a non-magnetic reference bead (Dynabeads). Bead positions were determined from images of the beads recorded at 120 Hz by a CMOS camera (Mikrotron EoSens) with GPU-assisted real-time particle tracking^42^. The applied forces on each bead were calibrated using power spectral density analysis^71^. For the TIRF measurements, the flow cell was illuminated through the objective in total internal reflection geometry using a 642nm laser (Omicron). The emitted fluorescence was separated using a dichroic mirror (R:633-643 nm/T:660-750 nm; Chroma). A laser rejection filter (642 nm; Chroma) removed residual laser light while a band-pass filter (A: 750-1100 nm; Semrock) blocked residual IR tracking light. The images were recorded with an EMCCD camera (Andor) at frame rates of 20 Hz (dwell time measurements) or 50 Hz (lateral DNA diffusion measurements). A frame rate of 10 Hz was used for the images shown in Figs. 1f; 2a, d; Supplementary Fig. 2. To generate trajectories of the emitted fluorescence of single bound Cascade complexes, the fluorescent spot in the acquired images was analyzed in Matlab. During the experiments, desired forces on the DNA construct could be set by placing the magnets at a particular distance from the flow cell according to the calibration results. Supercoiling of DNA was achieved by turning the magnets. Once a plectonemic superhelix is formed, the resulting torque depends mainly on the applied stretching force as well as the ionic strength of the solution. The torque was calculated based on previous theoretical work^72,73^. Time trajectories of the DNA length were recorded at 120 Hz and typically smoothed with a sliding average to 3 Hz for analysis. For enzyme measurements, St-Cascade was added in fluorescence buffer (20mM Tris-HCl pH 8.0, 150mM NaCl, 0.5 mg ml^-1^ BSA, 2mM Trolox, 5mM PCA, 75nM PCD) at a concentration of 1 nM or 2 nM. After adding St-Cascade, DNA length changes and fluorescence signals were monitored in real-time.

### Single-molecule diffusion measurements

Flow cell preparation was performed in the same manner as for the combined magnetic tweezers and fluorescence measurements. 15kbp DNA constructs without a matching target were bound on one side to 1µm diameter superparamagnetic beads (Dynabeads) and to the flow cell on the other. Instead of two cubical magnets, as for standard magnetic tweezers measurements, a cylindrical magnet was employed, exerting a lateral pulling force and stretching the DNA horizontally along the surface of the flow cell. Measurements were performed at a frame rate of 50 Hz. Enzymes were added to the flow cell in fluorescence buffer at a concentration of 1nM.

### Target recognition measurements

Target recognition measurements were performed in a custom-built magnetic tweezers setup^43^ at room temperature, as described before^17^. DNA molecules were bound with their biotin-labeled ends to streptavidin-coated magnetic beads with 0.5 µm diameter (Ademtech). A flowcell was covered with anti-digoxigenin and the DNA was flushed in, allowing tethering via the digoxigenin-labeled end. DNA length determination, force calibration and torque application were carried out as described for the combined magnetic tweezers and fluorescence measurements (see above). Wild-type St-Cascade (0.5 nM) was flushed into the flowcell in binding buffer (20 mM Tris-HCl pH 8.0, 150 mM NaCl, 0.1 mg ml^-1^ BSA). Negative supercoiling was applied to facilitate and observe R-loop formation. Once an R-loop was locked, identified by a constant DNA length corresponding to a full R-loop over several minutes, positive supercoiling (at a force of ∼2.5 pN) was applied to enforce dissociation of St-Cascade. The process was repeated after a subsequent R-loop was formed. Time trajectories were acquired with a CMOS camera at a frame rate of 120 Hz and smoothed to 3 Hz for analysis using a sliding average filter.

### Bulk binding assay

Matching or non-matching oligos (15nM) were incubated with wild-type or Cy5-labeled St-Cascade (30nM) for 1h in St-Cascade binding buffer (20 mM Tris-HCl pH 8.0, 150 mM NaCl). Binding was analyzed using an 8% native polyacrylamide gel with an acrylamide/bisacrylamide ratio of 29:1 and imaged using a ChemiDoc MP imaging system.

### Data analysis

Data analysis was carried out in Matlab (transition points in trajectories using Hidden Markov modeling) and Python (Maximum likelihood estimation). All plots were generated in Origin (OriginLab). Processing of the fluorescent images and generation of kymographs was carried out using custom-written software in Labview (National Instruments) and ImageJ^74^. Single-particle tracking of trajectories of individual Cascade complexes for mean-square-displacement analysis was carried using FIESTA^75^ Simulated magnetic tweezers and TIRF trajectories (Supplementary Methods 1, 3) were produced in LabView and Matlab, respectively. The functions used for maximum likelihood estimations are described in Supplementary Methods 2. The modeling of the target search and target recognition was performed in Python and the derivations are described in detail in Supplementary Methods 4-7.

