## Supplementary Information for "Dynamic interplay between target search and recognition for the Cascade surveillance complex of type I-E CRISPR-Cas systems"

### Supplementary Figures and Tables

**Table S1**. Oligonucleotides and plasmids used in the study.

| Oligo ID | Description | Sequence |
|---|---|---|
| TS89 | Cloning of Cse8e with C-terminal His-tag to pBAD24, forward Eco31I [NcoI tilt] | ggtctcagatccaatgagtcggtttaatttacttgatgaac |
| TS90 | Cloning of Cse8e with C-terminal His-tag to pBAD24, reverse XhoI | ctcgagttttaactgttgacgaagccaaaaatcg |
| IR76 | C252S and C262S mutagenesis of Cas8e, forward | gaatccggtaatgtagaaaatattcagcgaccttcctgggaacgtaaaag |
| IR77 | C252S and C262S mutagenesis of Cas8e, reverse | cccaggaaggtcgctgaatattttctacattaccggattcattatacg |
| IR92 | V76C mutagenesis of Cas6, forward | caaaagcttgaaaaatatggttgtgtaggaagcgctc |
| IR93 | V76C mutagenesis of Cas6, reverse | cctacacaaccatatttttcaagcttttgtaaattagg |
| JM_Not1_forward | Forward primer for making construct for combined magnetic tweezers and fluorescence measurements | tattacgccagcggccgcaaggggggatgtgctgcaag |
| PA_Cascade_Rev_SpeI_2.1 | Reverse primer for making construct for combined magnetic tweezers and fluorescence measurements | gatcagttgggtgcacgagtggactagtgtgaa |
| PS1-T20f/r | Targets with AAN-PAM with 20 PAM-distal mismatches cloned into pUC19 for combined magnetic tweezers and fluorescence measurements | gaccacccttttgatataatatacctatatcataccggagggtgcgtattcggcagatacgttctgagggaa |
| PS1-T20f/r-AG | Targets with AGN-PAM with 20 PAM-distal mismatches cloned into pUC19 for combined magnetic tweezers and fluorescence measurements | gaccacccttttgatatagtatacctatatcataccggagggtgcgtattcggcagatacgttctgagggaa |
| PS1-T20f/r-CC | Targets with CCN-PAM with 20 PAM-distal mismatches cloned into pUC19 for combined magnetic tweezers and fluorescence measurements | gaccacccttttgatatcctatacctatatcataccggagggtgcgtattcggcagatacgttctgagggaa |
| PS1-T1f | Targets with one PAM-distal mismatch for target recognition measurements | gaccacccttttgatataatatacctatatcaatggcctcccacgcataagggcagatacgttctgagggaa |
| PS1-T2f | Targets with two PAM-distal mismatches for target recognition measurements | gaccacccttttgatataatatacctatatcaatggcctcccacgcataacggcagatacgttctgagggaa |
| PS1-T4f | Targets with four PAM-distal mismatches for target recognition measurements | gaccacccttttgatataatatacctatatcaatggcctcccacgcatttcggcagatacgttctgagggaa |
| PI_ZT_PNL_F | Matching target oligo for EMSA | ggaccacgcataatatacctatatcaatggcctcccacgcataagcagtg |
| Nonmatching-F | Non-matching target oligo for EMSA | cggaccacccataagctgtctttcgctgctgagggtacg |
| **Plasmid** | **Description** | **Reference** |
| **pUC19** | Plasmid used for cloning plasmids with different CRISPR targets | New England Biolabs, #N3041S |
| pCRh | Contains homogeneous CRISPR region with 6 identical spacers | Sinkunas et al., 2013[1] |
| pCDF-Cascade-Cas8e-C252S-C262S-Cas6-V76C | Plasmid encoding double mutant of Cas8e C252S-C262S and a mutant of Cas6 V76C | This study |
| pBAD24-CHis | Used for cloning of Cas8e fusing with C-terminal $His_6$-tag | Thermo Fisher Scientific |
| pBAD-Cas7-C-His | Contains Cas7 protein with C-terminal $His_6$-tag under *araC* based promoter. | Sinkunas et al., 2013[1] |
| pCas3 | Contains Cas3 protein with C-terminal $His_6$-tag under *araC* based promoter. | Sinkunas et al., 2013[1] |
| pBAD-Cas8e-C-His | Contains Cas8e protein with C-terminal $His_6$-tag under *araC* based promoter. | This study |
| pSP1-AA | Specific target with PAM AA for combined magnetic tweezers and fluorescence measurements at low torque | Sinkunas et al., 2013[1] |
| pSP3-AA | Non-specific target with PAM AA | Sinkunas et al., 2013[1] |
| **Bacterial strains** | **Description** | **Reference** |
| *Escherichia coli* DH5a | Used for plasmid cloning, plasmid purification | Thermo Fisher Scientific, #18265017 |
| *Escherichia coli* BL21 (DE3) | Used for protein expression | New England Biolabs, #C2527 |

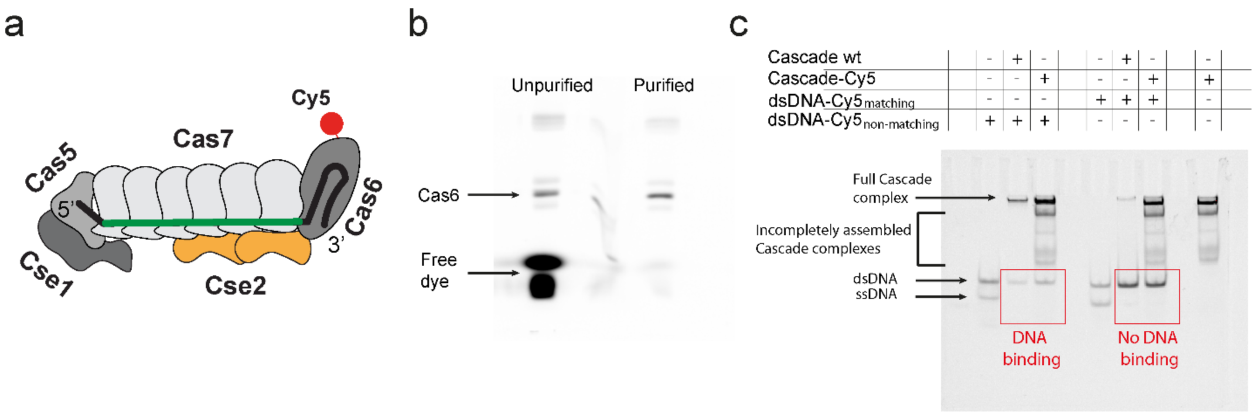

**Figure S1. Labeling St-Cascade with a Cy5 dye for TIRF measurements.** a Labeled St-Cascade complex. The complex is labeled via a malemeide linker on the Cas6 protein. b SDS-PAGE showing the successfully labeled Cas6 before and after purification  c Electrophoretic Mobility Shift Assay. St-Cascade labeled with Cy5 and wild-type binding behaviour on matching and non-matching dsDNA targets labeled with Cy5. Both St-Cascade variants bind to the matching target but not to the non-matching dsDNA target. Both variants show non-specific binding to ssDNA.

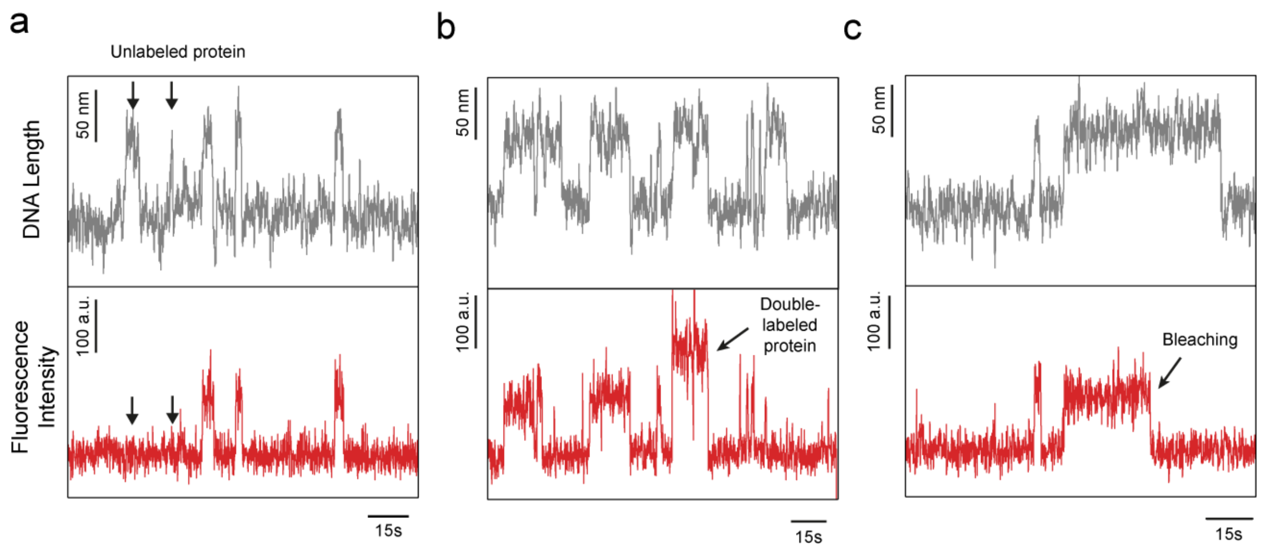

**Figure S2. Correlated trajectories magnetic tweezers and fluorescence trajectories monitoring unlabeled and dimeric Cascade complexes as well as fluorophore bleaching.** All shown trajectories were taken at 0.2pN and -6 turns ($\Gamma$ = -4.7 pN nm). The DNA length trajectories (grey) were taken at 120 Hz and smoothed to 3Hz while the fluorescence trajectories (red) were recorded at 10 Hz. **a** R-loop formation events without detectable binding events (indicated by black arrows) indicate the binding of unlabeled St-Cascade complexes. By counting the binding events of labeled and unlabled complexes, a labeling efficiency of St-Cascade of 65±10% was determined from the single-molecule experiments. **b** Occasional observation of binding events (7% of events) with doubled intensity (black arrow) indicated the binding of doubly labeled St-Cascade or a St-Cascade dimer. These events were considered in our quantitative analysis as they did not show an altered behavior **c** Occasional disappearance of the fluorescence signal before R-loop collapse was attributed to bleaching of the Cy5-fluorophore. In the analysis, these events were included for determining the delay between binding and R-loop formation but not for dissociation.

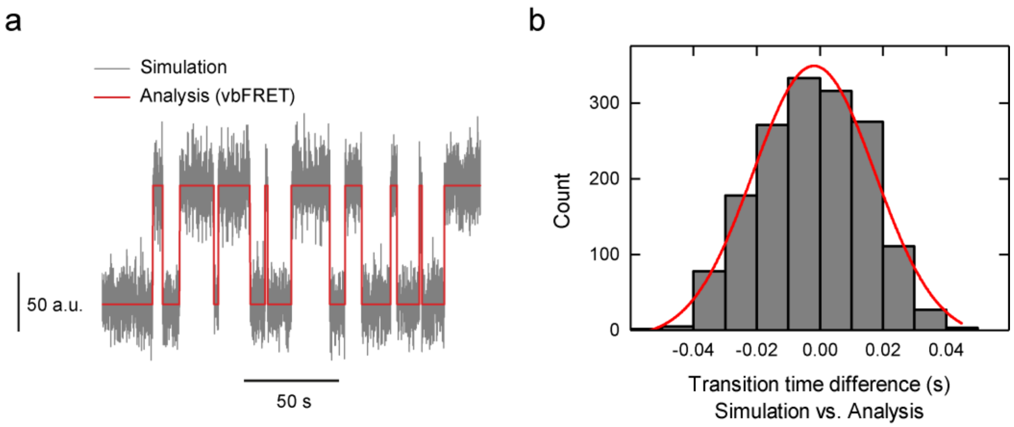

**Figure S3 Analysis of transition times in simulated fluorescence trajectories. a** Simulated fluorescence trajectories (grey) comprising multiple binding-dissociation events with an average length of 10 s. The two-state approximation of the trajectory from Hidden Markov modelling is shown in red. **b** Distribution of the error of individual transition times determined from the simulated trajectories (grey bars, $N = 1602$). A Gaussian distribution is depicted in red. It provided a mean of 1ms and a standard deviation of 17 ms.

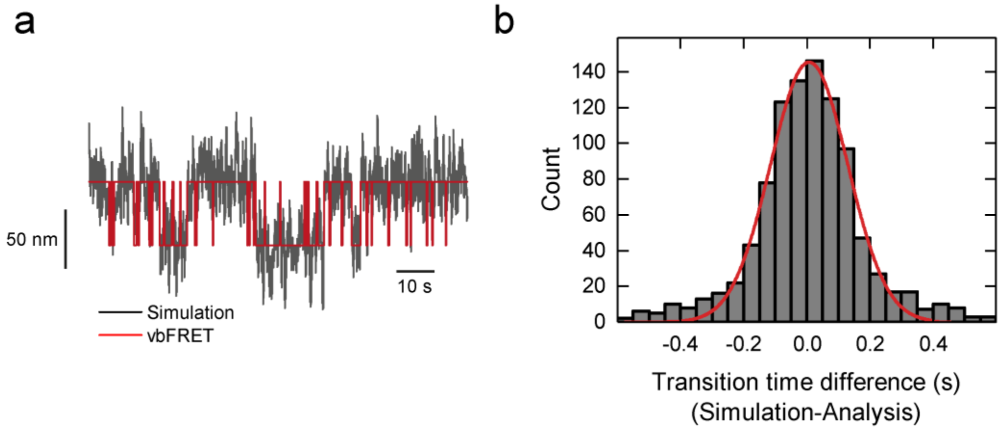

**Figure S4 Analysis of transition times in simulated magnetic tweezers trajectories. a** Simulated magnetic tweezers trajectory with multiple R-loop formation events (black) and a two state approximation of the trajectory from Hidden Markov modeling (red). **b** Difference between the nominal transition times from the simulated trajectory and the transition times found in the simulated trajectories. The histogram of the values is depicted in black, while the maximum likelihood function is depicted in red.

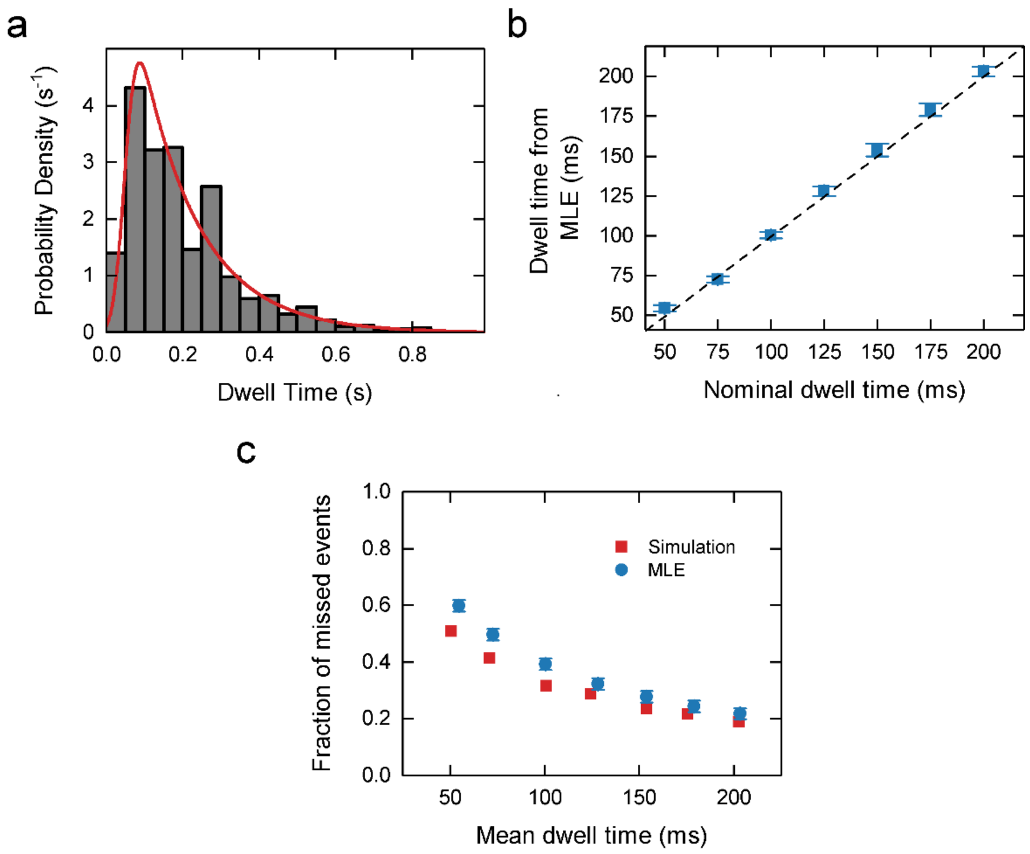

**Figure S5. Estimating the fraction of undetected short binding events by MLE. a** Dwell time histogram obtained from simulated fluorescence trajectories (50 ms integration time) using a nominal dwell time of 154 ms (grey bars, *N* = 1865). The result of the MLE analysis is shown as a red line providing $\tau = 154 \pm 4$ ms and $\mu = 53 \pm 10$ ms **b** Mean dwell times found by the MLE analysis compared to the nominal mean dwell times used in the simulations (blue squares). The dotted line corresponds to the identity between both axis values. **c** Fraction of missed events retrieved from the MLE analysis (blue circles) compared to the actual fraction of missed events by Hidden Markov modelling (red squares).

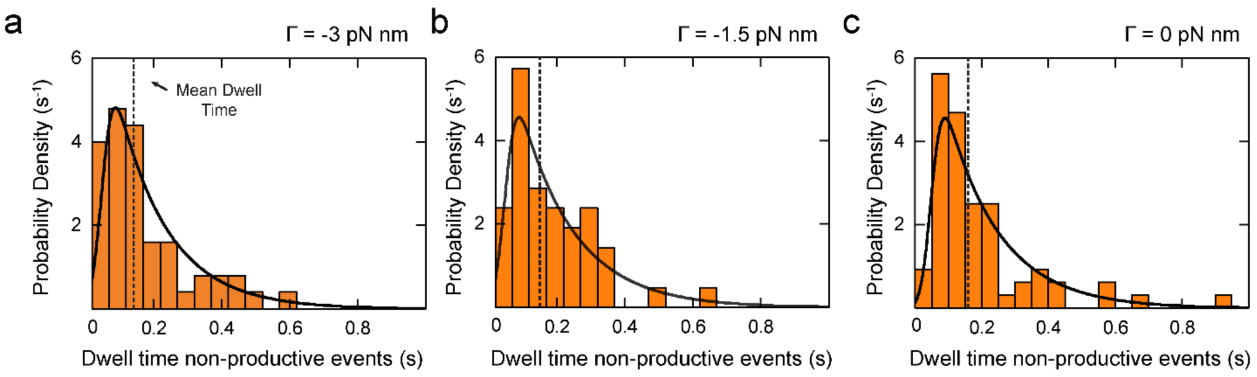

**Figure S6. Dwell time histograms of the non-productive binding events for different torques**. The black lines represent the maximum likelihood estimate of the data, while the dashed line indicates the mean dwell time. **a** Histogram for $\Gamma$ = -3 pN nm ($N =$ 50, $\tau_{mean}$ = 135 ± 19 ms). **b** Histogram for $\Gamma$ = -1.5 pN nm ($N =$ 50, $\tau_{mean}$ = 145 ± 23 ms). **c** Histogram for $\Gamma$ = 0 pN nm ($N =$ 114, $\tau_{mean}$ = 160 ± 17 ms).

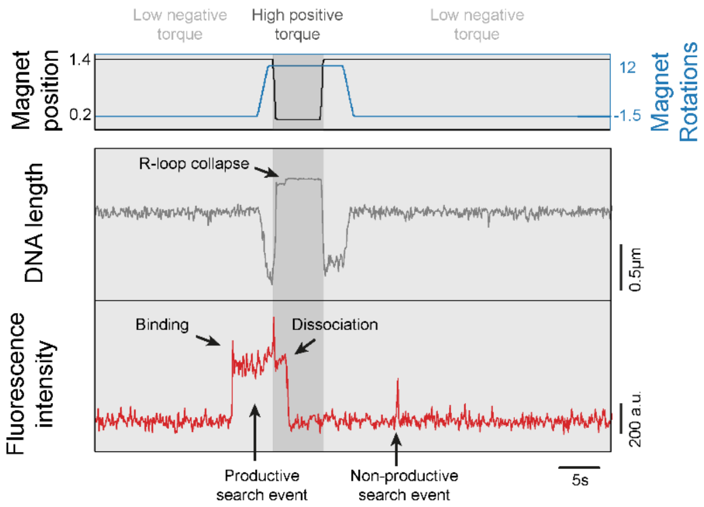

**Figure S7. Correlated DNA length and fluorescence trajectories with productive and non-productive search events at low torque (Γ= -1.5 pN nm, F = 0.2 pN).** The DNA length (grey) trajectory was recored at 120 Hz and smoothed to 10 Hz, while the fluorescence trajectory (red) was recorded at 20Hz. Long lasting binding events corresponded to productive target search events yielding a locked R-loop. Locked R-loop formation was verified and reversed by transiently applying a high positive torque (dark grey shaded area, Γ= 15 pN nm, F = 1.6 pN). This promoted R-loop collapse seen as a pronounced DNA length jump. In contrast, short binding events corresponded to non-productive search event without R-loop formation.

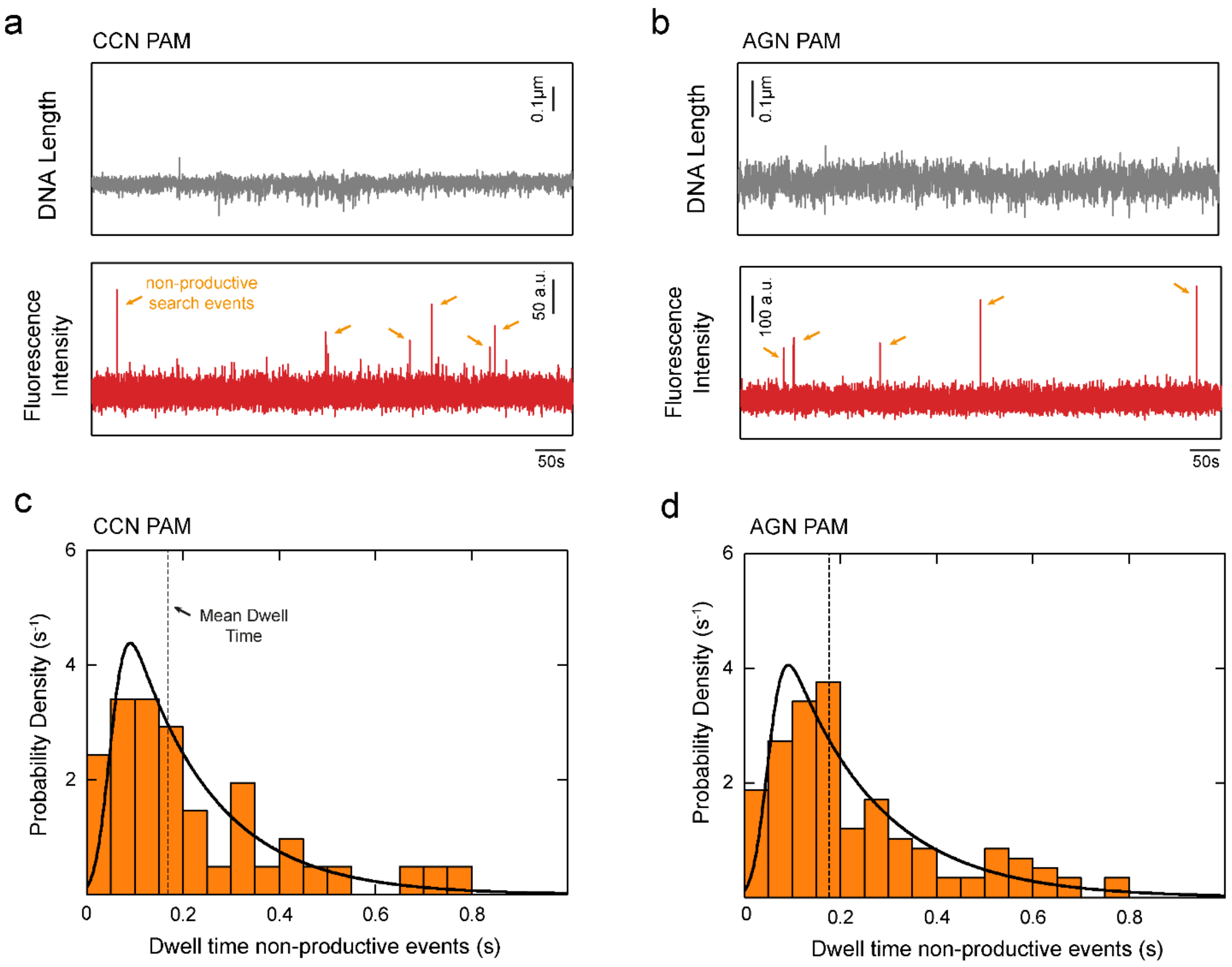

**Figure S8. Correlated DNA Length and fluorescence trajectories and dwell time histograms of the non-productive binding events for targets with non-cognate PAMs. a** Trajectories for the non-cognate CCN PAM recorded at $\Gamma$ = -4.7 pN nm (F=0.2 pN, -4 turns). The orange arrows point at non-productive search events. The DNA length was recorded at 120 Hz and smoothed to 3 Hz. The fluorescence signal was recorded at 20 Hz. **b** Trajectories for the non-cognate AGN PAM recorded at $\Gamma$ = -4.7 pN nm(F=0.2 pN, -5 turns). **c** Histogram for a CCN PAM ($N$ =41, $\tau_{mean}$ = 170 ± 30 ms). Black lines represent the maximum likelihood estimate of the data, while dashed lines indicate the mean dwell time. The data was taken at a torque of $\Gamma$ = -4.7 pN nm. **d** Histogram for a AGN PAM ($N$ = 119, $\tau_{mean}$ = 180 ± 20 ms). Black lines represent the maximum likelihood estimate of the data, while dashed lines indicate the mean dwell time. The data was taken at a torque of $\Gamma$ = -4.7 pN nm.

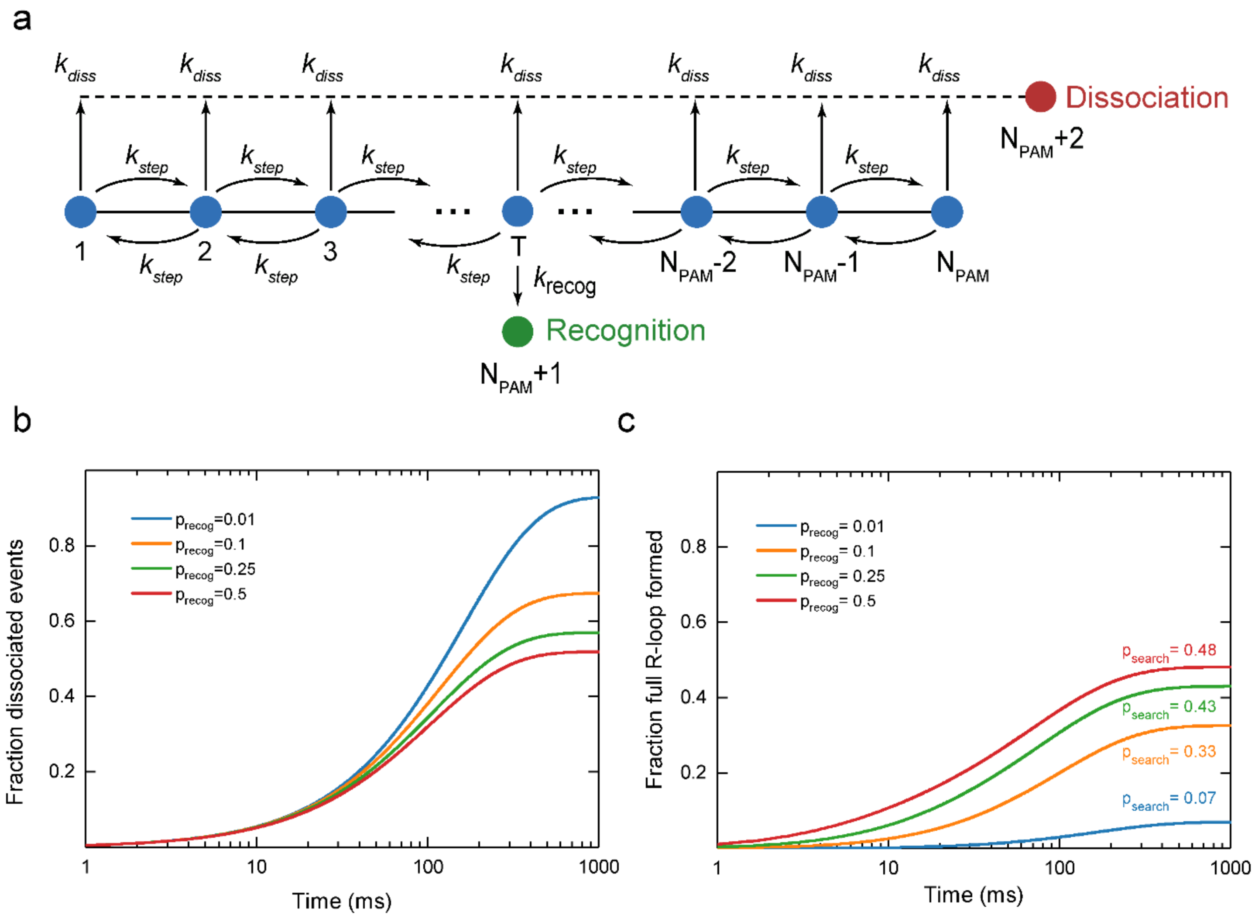

**Figure S9 Simulation of the target search model. a** The $N_{PAM}$ sequential states represent permissive PAMs with one of the PAMs ($T$) containing the adjacent target sequence. After random binding to any of the PAMs, Cascade stochastically transits to either of the two next-neighbor PAMs at equal rates $k$. At every PAM, Cascade can dissociate with rate $k_{diss}$, corresponding to an irreversible transition to State $N_{PAM}+2$. At the target PAM, $T$, Cascade can additionally recognize the target with rate $k_{recog}$, corresponding to an irreversible transition to state $N_{PAM}+1$. **b** Kinetics of non-productive target search events for which Cascade dissociated without recognizing the target. **c** Kinetics of productive target search events for which Cascade recognized the target. The model was solved for a lattice length $N=60$ and different target recognition probabilities per target binding $p_{recog}$. The fraction of productive events that is asymptotically approached at long times corresponds to the efficiency $p_{search}$ of the target recognition process.

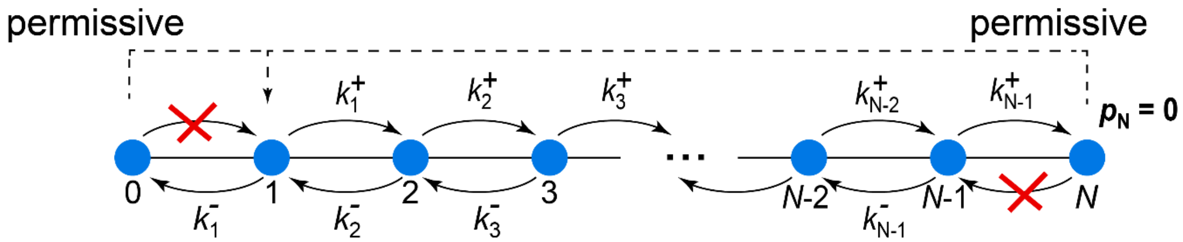

**Figure S10 Scheme of the one-dimensional random walk model for target recognition.** The probability to reach position $N$ when starting from position 1 without returning to zero is calculated. This problem can be treated by introducing permissive boundaries at position 0 and $N$. Particles that reach either of these positions are instantaneously placed back to position 1. The probabilities to reach either position $N$ or position 0 are proportional to the fluxes in either direction.

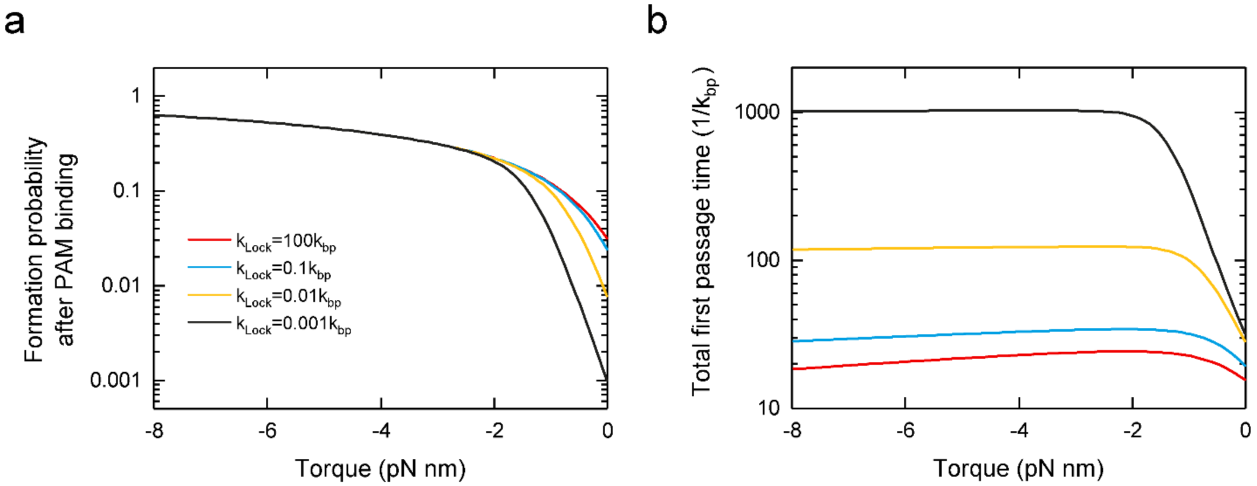

**Figure S11 Probability of locked R-loop formation as well as the mean duration of an R-loop formationevent as function of the bias of the energy landscape**. **a** Modeled formation probability after PAM binding as function of torque for different locking rates. **b** Total first passage time for R-loop formation function of torque. Shown are the numerical results for an R-loop length $N$ = 32 and various locking rates, where $k_{Lock}$ = 100$k_{bp}$ corresponds to the limit of very fast locking, i.e., negligible time.

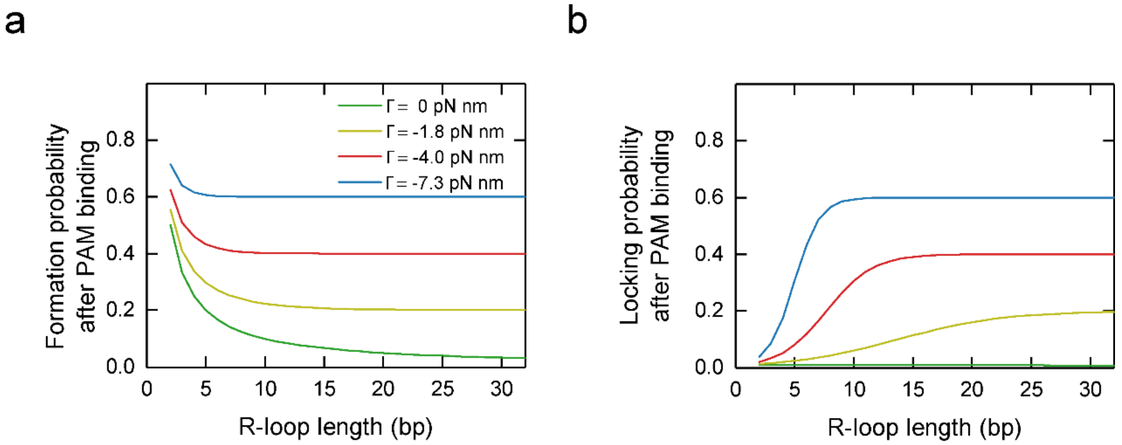

**Figure S12 Target recognition probability upon PAM binding as function of the R-loop length.** (Left) Formation probability of an unlocked R-loop vs. R-loop length plotted for differently biased energy landscapes. (Right) Probability for locked R-loop formation vs. R-loop length using a locking rate $k_{Lock}$= 0.01$k_{bp}$. The biases of the energy landscapes are colored as in the left plot.

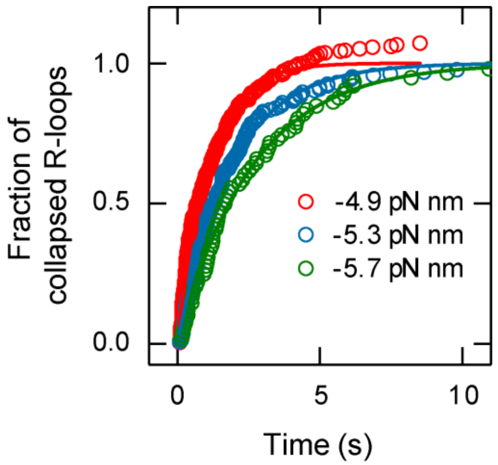

**Figure S13 Cummulative probability that a fully formed R-loop has collapsed to the intermediate state as function of time exemplary for two terminal mismatches.** Experimental results of individual collapse events for varying torques (-4.9 pN nm, red circles, N= 230; -5.3 pN nm, blue circles, N=160; -5.7 pN nm, green circles, N=68). Single exponential fits are depicted as lines in the respective colours.

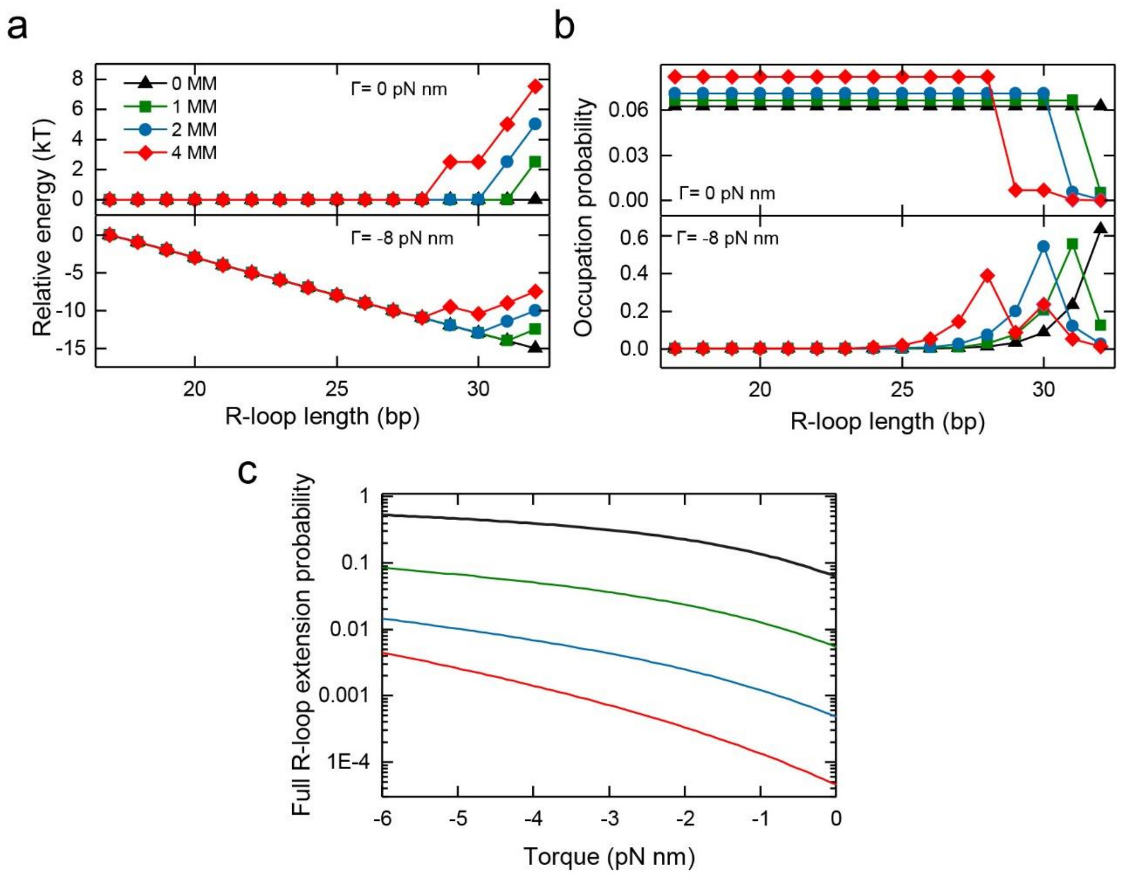

**Figure S14 Modeling the energy landscapes of R-loop formation by St-Cascade. a** Model energy landscapes of R-loop formation for different numbers of PAM-distal mismatches at zero torque (top) and at a torque of $\Gamma$ = -8 pN nm using a mismatch penalty of $\Delta G_{MM} = 2.5\ k_b T$. The depicted free energy is relative to the free energy at position 17. **b** Occupation probabilities of the R-loop length calculated for the different energy landscapes in **a** at zero torque and at a torque of $\Gamma$ = -8 pN nm. **c** Occupation probability of the full R-loop length, i.e. the probability that the R-loop extends to a length of 32 bp, as function of torque for different numbers of PAM-distal mismatches.

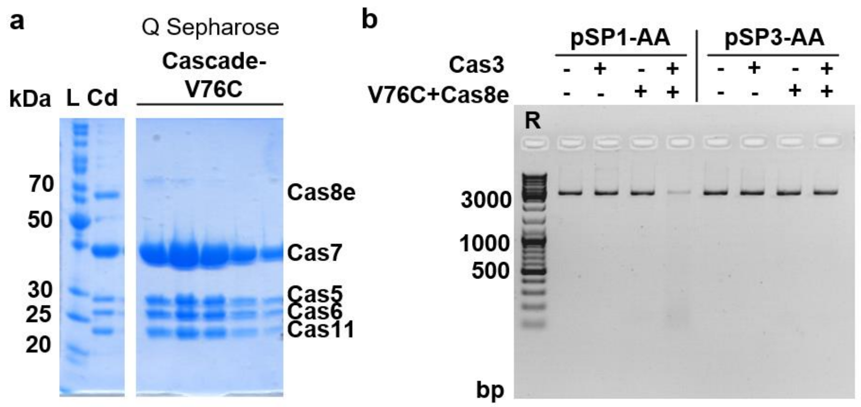

**Figure S15. Purification and cleavage assays of Cascade-Cas6-V76C effector complex.** a. 12 % SDS-PAGE after purification of Cascade through Q-Sepharose column. L represents PageRuler™ Unstained Protein Ladder, Cd – the wt Cascade effector complex as described in (Sinkunas et al., 2013). Next lines represent a peak of Cascade-Cas8e-C252S-C262S-Cas6-V76C complex (later on called Cascade-Cas6-V76C) on Q Sepharose column. Cas8e C252S-C262S mutant is missing from the complex. b. Cascade-Cas6-V76C activity assay with Cas3 nuclease-helicase. R stands for MassRuler DNA Ladder Mix. 3 nM pSP1-AA (specific target) or pSP3-AA (non-specific target), 100 nM Cas3, 100 nM Cascade and 100 nM Cas8e-C-His (or their buffers, respectively) incubated in the Nuclease buffer (10 mM Tris–HCl (pH 7.5), 75 mM NaCl, 40 mM KCl, 7% (v/v) glycerol, 1.5 mM MgCl2, 0.1 mM NiCl2, 2 mM ATP). The reaction was started by adding Cas3 to the mixture, incubated at 37°C and stopped after 40 min. Reactions were stopped by adding 1/3 of the reaction volume of “stop solution” containing 67.5 mM EDTA, 27% (v/v) glycerol, 0.3% (w/v) SDS and 0.1% w/v Orange G. The reaction products were visualized on 0.8 % agarose DNA gel and a voltage of 3 V/cm followed by visualization using ethidium bromide staining.

### Supplementary Methods

#### 1 Quantifying the accuracy of the transition point determination in fluorescence and magnetic tweezers trajectories

The determined dwell times connected to the target search process were in the range of the resolution limits of the magnetic tweezers assay and the fluorescence signal acquisition. Extraction of reliable dwell time estimates required an elaborate data analysis procedure. Both the fluorescence and the DNA length trajectories exhibited considerable noise that limited the accuracy at which transition points could be determined. To quantify the accuracy of the transition point determination, we used simulated fluorescence and DNA length trajectories including transitions and performed our data analysis in the same manner as for the experimental data. This allowed us to obtain realistic dwell time estimates as well as errors.

*Accuracy of transition points in fluorescence trajectories*

The accuracy of transition points in fluorescence trajectories was mainly limited by the Poisson distributed shot noise of the photon statistics[2] as well as the finite camera integration time of 50 ms. To simulate fluorescence trajectories, we first simulated two-state binding/dissociation trajectories obeying first order kinetics using $k_{bind}$= 0.05 $s^{-1}$ and $k_{diss}$= 0.1 $s^{-1}$ at a time resolution of 0.5 ms. We then split the trajectories into successive 50 ms windows and drew for each window random photon numbers from a Poisson distribution. The expected (mean) photon number $\lambda$ depended on the relative length of the unbound and bound states within each window. We assumed average photon numbers of $\lambda = 250$ and $\lambda = 350$ if the time window was fully occupied by the unbound state and the bound state, respectively. This provided a signal change of 100 photons upon binding, such that we obtained signal-to-noise ratios of $SNR \approx \Delta\lambda/\sqrt{\lambda} \approx 10$ as experimentally observed (Supplementary Fig. 3a).

We next analyzed the transition points in the simulated fluorescence trajectories in the same manner as the experimental trajectories using a Hidden Markov modelling algorithm[3]. The time error in determining the transition points was then calculated from the difference between input transition points for the simulations and output transitions from the analysis. The transition time errors were approximately Gaussian distributed with a mean of $\mu_{Fl} = 1.0 \pm 0.5\ \mathrm{ms}$ indicating that on average the positions of transition points were correctly determined. The standard deviation of the distribution, corresponding to the accuracy of a single measured transition time, was $\sigma_{Fl} = 17$ ms (Supplementary Fig. 3b).

*Accuracy of transition points in magnetic tweezers trajectories*

The accuracy of transition times in DNA length trajectories was mainly limited by the response time of the DNA tethered magnetic bead to sudden DNA length changes as well as thermal DNA length fluctuations. To simulate DNA length trajectories with R-loop formation-collapse events, we first simulated trajectories with sudden (R-loop) transitions between two fixed DNA lengths obeying first order kinetics with $k_{R,form}$= $k_{R,coll}$ = 0.05s$^{-1}$ at a frame rate of 120 Hz. The length difference between the two states was 60 nm as measured for experimentally observed transitions. To simulate DNA length fluctuations, we approximated our nanomechanical system as a sphere being attached to a linear spring with spring constant $k$ that is moving along the z coordinate with drag coefficient $\gamma$. The spring constant was determined from the mean-squared fluctuations of the DNA length in experimental trajectories of supercoiled DNA as $k = k_B T/\langle z^2 \rangle = 7.1 \cdot 10^{-4} \mathrm{pN\,nm}^{-1}$. The drag coefficient was then determined from the measured characteristic cut-off frequency $f_c = 12\,Hz$ of the frequency spectrum of the length fluctuations[4] as $\gamma = k/2\pi f_c = 9.4 \cdot 10^{-6}\ \mathrm{pN\,nm}^{-1}\,\mathrm{s}$. DNA length fluctuations including R-loop transitions were then modelled using Brownian dynamics simulations[4] in which the two-state R-loop trajectories provided the equilibrium position for the spring extension. (Supplementary Fig. 4a).

We next analyzed the transition times in the simulated trajectories using Hidden Markov modelling[3] and calculated their errors with respect to the nominal transition times. The transition time errors were Gaussian distributed with a pronounced non-zero mean of $\mu_{MT} =$ 13 ms and a standard deviation of $\sigma_{MT}$ =137 ms (Supplementary Fig. 4b). The measured time delay was due to the finite response time $\tau = \gamma/k =$ 13.2 ms which the nanomechanical system required to adapt to changes of the DNA length. For the collapse of the R-loop, a similar error distribution was obtained. In the DNA length trajectories, the vbFRET algorithm detected more transitions than were present. Excess transitions were removed manually, similar as for the experimental trajectories, for which the fluorescence signal provided a corrective.

### 2 Maximum Likelihood Estimation of short dwell times

The determined mean values of the dwell times $\tau_{R,form}$, $\tau_{R,coll}$ and $\tau_{non-prod}$ were smaller or on the order of the width of the experimentally determined dwell time distributions. To obtain accurate estimates of the mean dwell times we applied maximum likelihood estimation (MLE). Assuming that the target search follows first order kinetics, the actual dwell times are exponentially distributed around the mean dwell time $\tau$, which corresponds to the decay time of the exponential:

$$p_\tau(t) = \frac{1}{\tau}\exp\left(-\frac{t}{\tau}\right) \tag{S1}$$

Each experimentally determined dwell time was calculated from the difference of two transition times with their own Gaussian distributed statistical errors as determined above. The experimentally determined dwell times are then, according to the central limit theorem, Gaussian distributed around the actual values, whose mean and variance equal the sum of the means and variances of the individual error distributions, respectively. Correspondingly, the dwell time error distribution of the non-productive events had a mean of $\mu = 2\mu_{Fl} \approx 0\ ms$ and a variance of $\sigma^2 = 2\sigma_{Fl}^2 = (24\ ms)^2$ since it was calculated from two transitions in the fluorescence trajectories. The dwell time error distributions for the productive events had a mean of $\mu = \mu_{Fl} + \mu_{MT} \approx 13\ ms$ and a variance of $\sigma^2 = \sigma_{Fl}^2 + \sigma_{MT}^2 = (138\ ms)^2$. In base cases the dwell time error distributions are described by:

$$p_{error} = \frac{1}{\sigma\sqrt{2\pi}}\exp\left(-\frac{(t-\mu)^2}{2\sigma^2}\right) \tag{S2}$$

The error distributions shift and broaden the actual exponential dwell time distribution $p_\tau$ as observed in the experiments. The expected measured dwell time distribution $p_{\tau,exp}$ can be mathematically described as a convolution of $p_\tau$ with the corresponding error distribution $p_{error}$:

$$p_{\tau,meas} = p_\tau * p_{error} \tag{S3}$$

Inserting the distributions and solving the convolution integral provides:

$$\begin{aligned} p_{\tau,exp}(t) &= \int_0^\infty \frac{1}{\tau\sigma\sqrt{2\pi}}\exp\left(-\frac{(t-\mu-t')^2}{2\sigma^2}\right)\exp\left(-\frac{t'}{\tau}\right)dt' \\ &= \frac{1}{2\tau}e^{\frac{1}{2\tau}\left(2\mu+\frac{\sigma^2}{\tau}-2t\right)}\,\mathrm{erfc}\left(\frac{\mu+\frac{\sigma^2}{\tau}-t}{\sqrt{2}\sigma}\right), \end{aligned} \tag{S4}$$

where $\tau$ is the mean of the exponential distribution and $\mu$ and $\sigma$ are the mean and standard deviation of the of the Gaussian distributed errors, respectively. The probability of measuring a set of $n$ experimental dwell times $t_i$ is described by the likelihood function

$$L = \prod_i p_\tau(t_i) = \prod_i \frac{1}{2\tau}e^{\frac{1}{2\tau}\left(2\mu+\frac{\sigma^2}{\tau}-2t_i\right)}\,\mathrm{erfc}\left(\frac{\mu+\frac{\sigma^2}{\tau}-t_i}{\sqrt{2}\sigma}\right). \tag{S5}$$

Taking the logarithm of the likelihood function simplifies the product and yields a sum:

$$\ln(L) = \sum_i \ln\left[\frac{1}{2\tau} e^{\frac{1}{2\tau}\left(2\mu + \frac{\sigma^2}{\tau} - 2t_i\right)} \text{erfc}\left(\frac{\mu + \frac{\sigma^2}{\tau} - t_i}{\sqrt{2}\sigma}\right)\right]. \quad \text{(S6)}$$

To find the most likely mean dwell time $\tau$ describing the measured dwell time distribution, $\ln(L)$ was numerically maximized for the particular set of data points $t_i$ and error parameters $\mu$ and $\sigma$. The estimated standard errors of $\mu$ and $\sigma$ were derived from the inverted Hessian matrix of the optimized function, i.e. the covariance matrix[5].

### 3 Estimating the number of short non-detected binding events of St-Cascade to DNA

The average dwell times for non-productive binding of St-Cascade to DNA were on the order of 150 ms. Given a frame rate of 20 Hz in the measurements, a fraction of short-lived events remained undetected. To reliably determine the search efficiency, we used our MLE approach to estimate the number of undetected events. Assuming that only events with a dwell time larger than a cut-off $t_c$ are detected, the dwell time distribution of the detected events corresponds to an exponential distribution that is shifted by $t_c$ to larger times. Convolution with the error distribution would smoothen the sharp transition around $t_c$ and yield reduced event numbers at low times (Supplementary Fig. 5a), as also seen experimentally (Fig. 2e, main text). To obtain a nominal dwell time distribution, the shift by $t_c$ can be achieved by setting the mean of the Gaussian error distribution to $\mu = t_c$. It can be shown that the fraction of non-detected events among all binding events is then given by:

$$r_{missed} = 1 - e^{\frac{t_c}{\tau}}. \quad \text{(S7)}$$

where $\tau$ is the nominal mean dwell time. Using $\tau$ and $t_c$ $(\text{i.e.}\,\mu)$ as free parameters in our MLE approximation of the experimental dwell times allowed us to estimate $t_c$ and thus to correct for the fraction of non-detected non-productive binding events.

To validate this approach, we simulated fluorescence trajectories with short binding events using exponentially distributed dwell times of mean $\tau$ (see above as described above). Detection of the binding events in the simulated trajectories and analysis of the dwell times (see above) provided dwell time distributions with a significantly reduced number of events at low dwell times, which could be well approximated by the expected dwell time distribution with detection cutoff $t_c$ (Supplementary Fig. 5a). We carried out the simulations and the analysis for a range of different dwell times and verified that the analysis returned, within error, the input dwell times (Supplementary Fig. 5b). Most importantly, from the simulations we could also determine the actual number of events that were missed by the Hidden-Markov analysis of the trajectories and compare it with MLE result. This revealed that the MLE provided realistic fractions of missed events over the whole range of tested dwell times albeit with a slight overestimation (Supplementary Fig. 5c).

### 4 Target search model

Following the binding to a random PAM on the DNA, we modeled the target search by St-Cascade as a one-dimensional random walk with stepping rate $k_{step}$ from one PAM to the next adjacent PAM in either direction (Supplementary Fig. 9a). The 1D PAM lattice was limited to $N_{PAM}$ lattice points to mimic the limited illuminated DNA length. One of the PAMs (the target PAM, position 20, taken from the actual DNA sequence used in the experiments) was selected to contain the adjacent target. The mean-squared-displacement of such a discrete random walk is provided by:

$$\langle x^2 \rangle = 2k_{step}\Delta^2 t = 2Dt \qquad \text{(S8)}$$

with $\Delta=$ 2.7 nm being the mean distance between the PAMs. $\langle x^2 \rangle$ has to match the observed mean-square displacement, such that the stepping rate could be obtained from the experimentally determined diffusion coefficient as $k_{step} = \frac{D}{\Delta^2} = 1920\ s^{-1}$.

At each PAM, St-Cascade could furthermore dissociate from the DNA with dissociation rate $k_{diss}$ =5.7 $s^{-1}$ (Supplementary Fig. 9 and main text) yielding a non-productive search event. Upon arrival at the target PAM at position $T$, St-Cascade could form an R-loop at rate $k_{recog}$, yielding a productive event, which is related to the target recognition probability as $p_{recog} = \frac{k_{recog}}{k_{recog}+k_{diss}+2k_{step}}$.

Our model system can be seen as a Markov process with $N_{PAM} + 2$ states, where state $N_{PAM} + 1$ represents the productive R-loop state and state $N_{PAM} + 2$ the unproductive dissociated state. Let $\vec{p}(t)$ be the vector containing the occupation probabilities of all states at any time $t$, the kinetics of $\vec{p}(t)$ and thus the search process is then described by[6]:

$$\frac{d}{dt}\vec{p}(t) = \overline{\overline{K}}\,\vec{p}(t)\ , \qquad \text{(S9)}$$

where $\overline{\overline{K}}$ is the rate matrix containing the rate constants $k_{i,j}$ of all possible transitions between all states $i, j$ of the system. For the scheme in Figure S9a, the rate matrix has the following form:

$$
\bar{\bar{K}} =
\begin{array}{c}
\begin{array}{cccccccc}
1 & 2 & \dots & T & \dots & N_{PAM} & & \\
\end{array} \\
\begin{pmatrix}
-k_{step} - k_{diss} & k_{step} & \dots & 0 & \dots & 0 & 0 & 0 \\
k_{step} & -2k_{step} - k_{diss} & \dots & 0 & \dots & 0 & 0 & 0 \\
0 & k_{step} & \dots & 0 & \dots & 0 & 0 & 0 \\
\dots & \dots & \dots & \dots & \dots & \dots & \dots & \dots \\
0 & 0 & \dots & k_{step} & \dots & 0 & 0 & 0 \\
0 & 0 & \dots & -2k_{step} - k_{diss} - k_{recog} & \dots & 0 & 0 & 0 \\
0 & 0 & \dots & k_{step} & \dots & 0 & 0 & 0 \\
\dots & \dots & \dots & \dots & \dots & \dots & \dots & \dots \\
0 & 0 & \dots & 0 & \dots & k_{step} & 0 & 0 \\
0 & 0 & \dots & 0 & \dots & -k_{step} - k_{diss} & 0 & 0 \\
0 & 0 & \dots & k_{recog} & \dots & 0 & 0 & 0 \\
k_{diss} & k_{diss} & \dots & k_{diss} & \dots & k_{diss} & 0 & 0
\end{pmatrix}
\end{array}
\begin{array}{c}
1 \\ 2 \\ 3 \\ \dots \\ T-1 \\ T \\ T+1 \\ \dots \\ N_{PAM}-1 \\ N_{PAM} \\ N_{PAM}+1 \\ N_{PAM}+2
\end{array}
\tag{S10}
$$

Eq. S9 is solved by

$$\vec{p}(t) = \exp\left(\bar{\bar{K}}t\right)\vec{p}(0) \tag{S11}$$

with $\exp\left(\bar{\bar{K}}t\right)$ being the matrix exponential of $\bar{\bar{K}}$ and $\vec{p}(0)$ being the initial probability distribution at $t = 0$. The exponential term can be expressed as

$$\exp\left(\bar{\bar{K}}t\right) = \bar{\bar{x}} \cdot \exp\left(\bar{\bar{\Lambda}}t\right) \cdot \bar{\bar{x}}^{-1} \ , \tag{S12}$$

where $\bar{\bar{x}}$ is the eigenvector matrix of $\bar{\bar{K}}$, and $\exp\left(\bar{\bar{\Lambda}}t\right)$ is a diagonal matrix, containing exponential decays in time with the eigenvalues $\lambda_i$ as decay constants:

$$\exp\left(\bar{\bar{\Lambda}}t\right) = \begin{pmatrix} \exp(\lambda_1 t) & 0 & \dots \\ 0 & \exp(\lambda_2 t) & \dots \\ \dots & \dots & \dots \end{pmatrix}. \tag{S13}$$

The entries of the matrix exponential are, according to Eq. S12, sums of the $N + 2$ exponential decays:

$$\left(\bar{\bar{x}} \cdot \exp\left(\bar{\bar{\Lambda}}t\right) \cdot \bar{\bar{x}}^{-1}\right)_{i,j} = \sum_{k=1}^{N_{PAM}+2} x_{i,k} x_{k,j}^{-1} \exp(-\lambda_k t). \tag{S14}$$

Combining Eqns. S11, S12 and S14, provides then for the occupation probability of state $i$ the following sum of the $N_{PAM}$+2 exponential decays:

$$p_i(t) = \sum_{k=1}^{N_{PAM}+2} \exp(-\lambda_k t) \sum_{j=1}^{N_{PAM}+2} p_j(0)\, A_{ijk} \tag{S15}$$

with $A_{ijk} = x_{i,k} x_{k,j}^{-1}$. For an initial binding of Cascade to a random PAM, the starting probabilities are given by:

$$\vec{p}(0) = (1/N_{PAM}, 1/N_{PAM}, \dots, 1/N_{PAM}, 0{,}0) \tag{S16}$$

With this solution and experimentally determined values for $k_{step}$ and $k_{diss}$, one can calculate the kinetics of productive ($p_{N+1}(t)$) and non-productive ($p_{N+2}(t)$) target recognition events for any set the lattice length $N_{PAM}$ and the target recognition efficiency $p_{recog}$ (Supplementary Fig. 9a). We validated the obtained kinetics using stochastic simulations of the random walk process. The fraction of productive events that is asymptotically approached at long times provides the efficiency $p_{search}$ of the target recognition process (Supplementary Fig. 9c). $p_{search}$was increasing with increasing $p_{recog}$ and generally larger than $p_{recog}$ due to multiple target site revisits during the search. From the kinetics, we furthermore calculated the mean event durations, i.e. the dwell times for non-productive and productive search events:

$$\tau_{R,form/non-prod} = \langle t_i \rangle = \sum_{k=1}^{N_{PAM}+2} 1/\lambda_k \sum_{j=1}^{N_{PAM}+2} p_j(0)\, A_{ijk} \qquad \text{(S17)}$$

with $i$ being either $N_{PAM}+1$ or $N_{PAM}+2$, respectively.

Dwell times for R-loop collapse were obtained from the calculated kinetics of non-productive events that started at the target site at position $T$ for which the initial occupancies changed to $\vec{p}(0) = (0, \dots, 0,1,0, \dots, 0)$. The number of target site revisits for productive events was obtained from 10,000 stochastic simulations of the target search process following the scheme in Supplementary Fig. 9a. The search process was always started at the target site and the number of revisits was counted for each productive search event.

### 5 Target recognition model

*Probability to finish a 1D random walk once the first step was taken*

Due to the reversibility of the R-loop formation process, an R-loop of a CRISPR-Cas effector complex that became nucleated upon PAM binding has, even on a perfectly matching sequence, a certain chance to collapse without full R-loop formation or locking. To provide a theoretical description for the probability $p_{recog}$ at which a matching target becomes recognized upon PAM binding, we model R-loop formation as a one-dimensional random walk process over $N + 1$ positions (0,1,2,...,$N$), where $N$ corresponds to the number of base pairs of the full R-loop (Supplementary Fig. 10). Within the model, position 0 corresponds to the unbound state, position 1 to the PAM-bound state with the first base-pair of the R-loop being formed and the positions from 2 to $N$ to all subsequent R-loop lengths. To include R-loop locking, an additional position $N + 1$ can be introduced. Transitions are only allowed between neighboring states. They are described by position-dependent rate constants $k_i^+$ for forward steps and $k_i^-$ for backward steps with $i$ indicating the position. Within the random walk model, the target recognition following PAM binding translates into a start of the walk at position 1. The walk is terminated when either Position 0 (non-successful recognition) or position $N$ (successful recognition) are reached. The target recognition probability is then the probability $p_{pass}$ for a successful termination of the walk. To obtain this probability but also the time scale of the walk in the frame work of the model, one introduces permissive boundaries at positions 0 and $N$ and places a single particle into the system.

Once the particle reaches a boundary, the particle is instantaneously placed to the start position 1. To obtain average quantities over many single-particle trajectories, we assume steady state conditions for this system, i.e. probabilities $p_i$ for the particle to be found at position $i$ that are time-invariant. Given the permissive boundaries we directly get $p_0 = p_N = 0$. In steady state, this results in a constant backward flux $j_-$ of particles from position 1 to 0 and a constant forward flux $j_+$ of particles from position 1 to $N$. The fluxes correspond to the rates of particles arriving at either boundary, i.e. to the rates of successful vs. unsuccessful recognition. The probability $p_{pass}$ to reach position $N$ is then provided by the forward flux divided by the total flux at position 1:

$$p_{pass} = \frac{j_+}{j_+ + j_-} = \frac{1}{1 + \frac{j_+}{j_-}}. \qquad \text{(S18)}$$

For the forward/backward flux $j_i^\pm$ between two neighboring positions $i$ and $i + 1$ we can write:

$$j_i^\pm = \pm k_i^+ p_i \mp k_{i+1}^- p_{i+1} \qquad \text{(S19)}$$

In steady state, the probability at any position $i$ does not change in time such that for a pure linear chain we have:

$$\frac{dp_i}{dt} = j_{i+1} - j_i = 0 \tag{S20}$$

Recursively, it follows that $j_i = j_0$ for all $i$ of a linear chain segment not containing branches. For the backward flux $j_-$, we then get from $p_0 = 0$ and Eq. S19:

$$\frac{p_1}{j_-} = \frac{1}{k_1^-} \tag{S21}$$

For the forward flux, we get from Eq. S19:

$$\frac{p_i}{j_+} = \frac{1}{k_i^+} + \frac{k_{i+1}^-}{k_i^+}\frac{p_{i+1}}{j_+} \tag{S22}$$

Thus, with the boundary condition $p_N = 0$, we can obtain all $p_i/j_+$ from $N$-1 to 1 in a recursive manner:

$$\frac{p_{N-1}}{j_+} = \frac{1}{k_{N-1}^+} \ ,$$

$$\frac{p_{N-2}}{j_+} = \frac{1}{k_{N-2}^+} + \frac{1}{k_{N-2}^+}\frac{k_{N-1}^-}{k_{N-1}^+} \ ,$$

$$\frac{p_{N-3}}{j_+} = \frac{1}{k_{N-3}^+} + \frac{1}{k_{N-3}^+}\frac{k_{N-2}^-}{k_{N-2}^+} + \frac{k_{N-2}^-}{k_{N-2}^+}\frac{k_{N-1}^-}{k_{N-1}^+}, \ldots$$

Since the probability distribution $\{p_i\}$ is normalized (single particle in the system) and $p_0 = 0$, one can easily derive an expression for the forward flux:

$$\sum_{i=1}^{N}\frac{p_i}{j_+} = \frac{1}{j_+}\sum_{i=0}^{N} p_i = \frac{1}{j_+} \tag{S23}$$

The local probabilities are then obtained by multiplying the derived values $p_i/j_+$ with $j_+$. From $p_1$ and Equation S21, one obtains $j_- = k_1^- p_1$. Using the principle of detailed balance, one can express the ratio of the rate constants between positions $i$ and $i+1$ directly as

$$k_{i+1}^-/k_i^+ = \exp(\Delta G_i/k_B T). \tag{S24}$$

The variable $\Delta G_i = G_{i+1} - G_i$ is the local bias of the energy landscape, i.e. the free energy difference between positions $i+1$ and $i$. With this, Eq. S22 becomes:

$$\frac{p_i}{j_+} = \frac{1}{k_i^+} + \frac{p_{i+1}}{j_+}e^{\Delta G_i/k_B T}. \tag{S25}$$

In a recursive manner we can obtain all $p_i/j_+$ as above, for which we obtain the following form final form:

$$\frac{p_{N-i}}{j_+} = \sum_{j=1}^{i} \frac{1}{k_{N-j}^+} \, e^{\left(G_{N-j} - G_{N-i}\right)/k_B T} \quad . \tag{S26}$$

Using this expression, we obtain $p_1/j_+$ and obtain after transformation:

$$j_+ = \frac{p_1}{\sum_{i=1}^{N-1} \frac{1}{k_{N-i}^+} \exp(G_{N-i} - G_1)/k_B T} = \frac{p_1}{\sum_{i=1}^{N-1} \frac{1}{k_i^+} \exp(G_i - G_1)/k_B T} \quad . \tag{S27}$$

With $j_- = k_1^- p_1$, we get for the success probability of the random walk:

$$p_{pass} = \frac{1}{1 + k_1^- \sum_{i=1}^{N-1} \frac{1}{k_i^+} \exp \frac{G_i - G_1}{k_B T}} . \tag{S28}$$

*Formation probability of a matching R-loop of N base pairs under constant bias*

To derive an expression for the formation probability of an (unlocked) R-loop of length $N$ following PAM binding, we assume a constant bias for the energy landscape for all R-loop states such that $G_{i+1} - G_i = E_{bias}$ for $1 \leq i < N$. The energy difference between position $i$ and 1 is given as $G_i - G_1 = (i-1)E_{bias}$. Furthermore, we assume position-independent values for the forward and backward stepping rates. Simplifying the exponential terms of the bias energy in Eq. S28 with the rate ratio between the backward and the forward stepping rate

$$s = \frac{k^-}{k^+} = e^{E_{bias}/k_B T} \tag{S29}$$

yields for the R-loop formation probability:

$$p_{Rloop} = \frac{1}{1 + e^{E_{bias}/k_B T} \sum_{i=1}^{N-1} e^{(i-1) \cdot E_{bias}/k_B T}} = \frac{1}{1 + \sum_{i=1}^{N-1} s^i} \tag{S30}$$

Using the sum formula of the geometric series, one gets

$$p_{Rloop} = \frac{1}{1 + s \frac{s^{N-1} - 1}{s - 1}} = \frac{s - 1}{s^N - 1} \tag{S31}$$

*Locked R-loop formation under constant bias*

We further consider ‘locking’ to derive a more realistic target recognition probability that includes the effects of the conformational changes during locking, which are slow compared to the single base pair steps. To this end, we introduce an additional locking step from position $N$ to $N + 1$ with the rate constant $k_N^+ = k_L$. The target recognition probability then becomes

$$p_{recog} = \frac{1}{1 + k^- \sum_{i=1}^{N} \frac{1}{k_i^+} e^{(i-1)\, E_{bias}/k_B T}} = \frac{1}{1 + \sum_{i=1}^{N-1} s^i + \frac{k^+}{k_L} s^N} \quad \text{(S32)}$$

This can be further transformed to:

$$p_{recog} = \frac{1}{1 + s \cdot \frac{s^{N-1} - 1}{s - 1} + \frac{k^+}{k_L} s^N} = \frac{s - 1}{s^N \left[1 + (s - 1)\left(\frac{k^+}{k_L}\right)\right] - 1} \quad \text{(S33)}$$

Assuming that the activation barriers for single base pair steps of R-loop formation are centered between subsequent R-loop positions, the bias-dependent forward and the backward stepping rates for R-loop formation can be expressed using Arrhenius-like terms:

$$k^{\pm} = k_{bp}\, e^{\mp\frac{E_{bias}}{2}} = k_{bp}\, s^{\mp\frac{1}{2}} \quad \text{(S34)}$$

with $k_{bp}$ being the forward and backward stepping rate in absence of bias. When plotting the target recognition probability as function of the (negative) bias of the energy landscape as applied by the supercoiling, one sees that it strongly increases with the applied bias, similar to the first passage time of R-loop formation (Supplementary Fig. 11). Reducing the locking rate reduces the target recognition probability only for weak but not for strong biases (Supplementary Fig. 11a). This can be better understood by looking at the probabilities of (unlocked) R-loop formation and locked R-loop formation as function of the R-loop length (Supplementary Fig. 12). For an unlocked R-loop formation we have high probabilities to form short R-loops due to the limited length of the random walk which drop with increasing R-loop lengths (Supplementary Fig. 12a). It is important to note that even for a mild downhill bias, the R-loop formation probability saturates at a minimum value already at short R-loop lengths. This means that the decision for formation or collapse of an R-loop is taken at the beginning of the random walk. Once the R-loop has been further expanded, it will for the given bias practically always form but not collapse anymore. This changes somewhat if a considerably slower locking step is added. Locked R-loop formation is particularly disfavored at short lengths, since the slow step represents a high kinetic (activation) barrier that needs to be overcome. The probability then increases with length in presence of a negative bias, reaching the same saturation values as for unlocked R-loops (Supplementary Fig. 12b). Due to the negative bias, the unlocked R-loops are already sufficiently stabilized compared to the activation barrier for locking. Locking is then just an additional step after R-loop formation and occurs without an additional collapse. By contrast, at zero or mild bias, the locking barrier still represents an additional hurdle that can promote R-loop collapse such that the locked R-loop formation is less likely than unlocked R-loop formation.

### 6 Determination of the apparent locking rate from magnetic tweezers trajectories

In the following we derive a simple formula to calculate the apparent locking rate from the magnetic tweezers trajectory in Fig. 5, main text. When the R-loop is in the full state $F$, R-loop collapse to the intermediate state $I$ and locking state $L$ are competing:

$$I \xleftarrow{k_I} F \xrightarrow{k_{lock,app}} L \tag{S35}$$

This kinetic scheme can be described by the following set of differential equations:

$$\begin{aligned} \frac{dp_F}{dt} &= -k_I p_F - k_L p_F \\ \frac{dp_I}{dt} &= k_I p_F \\ \frac{dp_L}{dt} &= k_L p_F \ , \end{aligned} \tag{S36}$$

where $p_F$, $p_I$ and $p_L$ are the probabilities that the R-loop is in the full, intermediate and locked state respectively. At $t = 0$, i.e. when the full state has just formed, we have the following starting conditions:

$$\begin{aligned} p_F(0) &= 1 \\ p_I(0) = p_L(0) &= 0 \end{aligned} \tag{S37}$$

The first differential equation is a simple first order decay, such that we get with the starting conditions:

$$p_F(t) = e^{-(k_I+k_L)t} \tag{S38}$$

Inserting into the second equation gives:

$$\frac{dp_I}{dt} = k_I e^{-(k_I+k_L)t} \tag{S39}$$

Integration gives for the time-dependent probability $p_I$ that a full R-loop has collapsed to the intermediate state:

$$p_I(t) = p_I(0) + \int_0^t k_I e^{-(k_I+k_L)t} dt = \frac{k_I}{k_I + k_L}\left(1 - e^{-(k_I+k_L)t}\right) \tag{S40}$$

as the only measurable probability. The probability $p_I(t)$ rises only to the amplitude:

$$\underbrace{\frac{k_I}{k_I + k_L}}_{p_{coll}} = 1 - \underbrace{\frac{k_L}{k_I + k_L}}_{p_{lock}} \tag{S42}$$

With $p_{coll}$ or $p_{lock}$ being the probabilities that a formed full R-loop collapses or becomes locked, respectively. The collapse occurs with the rate constant

$$k_{coll} = k_I + k_L \tag{S43}$$

If one determines $p_{lock}$ from the fraction of events that successfully get locked and $k_{coll}$ from fitting the probability of collapse for fully formed R-loop (Supplementary Fig. 13) using Eq. S40, one can calculate $k_I$ and $k_L$ from:

$$\begin{aligned} k_I &= (1 - p_{lock})k_{coll} \\ k_L &= p_{lock}\, k_{coll} \end{aligned} \tag{S44}$$

### 7 Estimation of the 'real' locking rate

Due to their reversibility, unlocked R-loops do not possess a fixed length but rather sample different lengths dependent on the energy landscape of R-loop formation. The results of our experiments suggested (see main text) that R-loop locking is only triggered if the R-loop extends over its full length. Given that the unlocked R-loop samples the full length only for a fraction of the time, the measured torque-dependent apparent locking rate would be considerably lower than the real locking rate $k_{lock}$, i.e., the rate at which the locking transition occurs, once a full R-loop has been formed. The relation between both rates would be given as:

$$k_{lock,app} = p_{32}\, k_{lock}\ , \qquad \text{(S45)}$$

where $p_{32}$ is the probability that the R-loop extends all the way to the last base pair. The probabilities $p_i$ that an R-loop extends over $i$ base pairs can be determined from the energy landscape of R-loop formation using the Boltzmann distribution:

$$p_i = \frac{1}{Z} e^{-G_i/k_B T} \qquad \text{with} \qquad Z = \sum_{i=1}^{32} e^{-G_i/k_B T} \qquad \text{(S46)}$$

being the partition function $Z$ that ensures normalization of the probabilities for all states. As described above, we assumed an energy landscape with a constant negative bias from the applied supercoiling. For each PAM-distal mismatch, a (positive) free energy penalty was introduced at its position[7,8] since disrupted base pairing is not compensated by base pairing between guide RNA and DNA target strand. No penalty was introduced for a target mutation at position 30 due to the base flips at this position observed in the Cascade structure[9]. Resulting model energy landscapes are shown in Supplementary Fig. 14a for different numbers of terminal mismatches and applied negative torque. Elevated negative torque increases the occupancy of the full R-loops state as it becomes energetically favored, while PAM distal penalties decrease the occupancy of the full R-loop state drastically (Supplementary Fig. 14b, c). Finally, using the simplified model for $k_{lock,app}$, the measured apparent locking rate data could be globally fitted using $k_{lock}$ and a base-pair independent mismatch penalty as fit parameters (Fig. 5, main text). Agreement between data and model supported the idea that R-loop locking only occurs for a fully extended R-loop and that the torque dependence of the $k_{lock,app}$ is mainly due to an increasing confinement of the R-loop to its full length with increasing negative supercoiling.
